# WRN deficiency generates oncogenic rearrangements at TA-rich tandem repeats

**DOI:** 10.64898/2026.10.16.758976

**Authors:** Meng Wang, Hui Pan, Anastasia Samarkina, Austin John Escobar, Noel Cruz Pacheco, Satoru Mizuhashi, Jamie D. Aquino, Katy K. Tsai, Adil I. Daud, Ivan EI-Sayed, Haruto Nishida, Sakuhei Fujiwara, Takashi Hirano, Sine Phelan, Allan O’Keefe, Sean Dinneen, A. Hunter Shain, Makoto Goto, Iwei Yeh, Boris C. Bastian

## Abstract

Werner syndrome, a premature-aging disorder caused by inherited loss of the WRN helicase, predisposes to a narrow set of cancers, including melanoma and sarcoma, through mechanisms that have been unclear. WRN resolves the secondary structures that TA-dinucleotide repeats form, a role established for the expanded repeats of mismatch-repair–deficient cancers. We show that WRN loss also destabilizes native, unexpanded TA-rich tandem repeats, which become fragile sites where chromosomes break and rearrange. In melanoma these rearrangements recurrently produce an identical 15.7Mb-long deletion that transcriptionally activates *TERT*, the earliest driver of these tumors, and separately disrupt tumor suppressors such as *CDKN2A/B* and *NF1*. The same class of rearrangement occurs in WRN-deficient cancers of other types. WRN deficiency is thus a source of instability at TA-rich tandem repeats that generates oncogenic rearrangements.

## Main Text

Werner syndrome (WS) is an autosomal recessive disorder of premature aging with a distinctive predisposition to a narrow set of rare cancers — sarcomas and the acral and mucosal subtypes of melanoma — tumors defined by clustered structural rearrangements rather than by the point-mutation signatures of environmental mutagens (*1–4*). WS is caused by biallelic loss of *WRN*, which encodes a RecQ-family helicase that resolves secondary DNA structures such as forks, bubbles, and G-quadruplexes during replication and repair (*5–7*) and stabilizes the G-rich lagging strand at telomeres (*8*). Why the loss of this activity predisposes so specifically to these cancers, and through what mutational mechanism, has remained unclear.

A clue comes from microsatellite-unstable (MSI) cancers, in which mismatch-repair deficiency leads to progressive expansion of TA-dinucleotide repeats (*9*). These expanded tracts form secondary structures that normally require *WRN* to resolve; when *WRN* is absent, they collapse into double-strand breaks, making *WRN* loss lethal in this setting (*9–13*). This established the view that *WRN* deficiency becomes consequential specifically at microsatellites *expanded* by mismatch-repair failure. Here we find that *WRN* loss has structural consequences at TA-rich tandem repeat sequences even without expansion: in microsatellite-stable, *WRN*-deficient cancers, these sequences become fragile sites at which chromosomes break and rearrange. In melanoma, these rearrangements recurrently activate *TERT* and disrupt tumor suppressors, driving the disease, and the same class of rearrangement occurs in *WRN*-deficient cancers of other types.

## Results

We performed Illumina short-read whole-genome sequencing (WGS) of four different acral and mucosal melanomas and corresponding normal tissue from three WS patients (WS1, WS2 and WS3) **(Fig. S1-S3, Table S1,** and **Data S1-S2)**. WS1 and WS2 were a pair of Japanese sisters (*14*, *15*), both harboring two heterozygous germline *WRN* nonsense mutations, R369* and R1305*, while WS3 was an unrelated case (*16*) who harbored a homozygous germline *WRN* nonsense mutation R1321*. From the sequencing data, we confirmed that the two melanomas from the left and right nasal cavities of WS1 were phylogenetically unrelated and harbored distinct sets of driver genes (**Fig S1**).

### A 15.7 Mb-long deletion upstream of *TERT* occurs exclusively in WRN-null melanomas

The four different melanomas from the three patients shared an identical 15.7 Mb-long DNA copy loss on chromosome 5p (**Fig. 1A**), which was absent in their respective germline DNAs. The breakpoints of the telomeric side (BP_tel_) of the region of DNA copy loss were located 12 kb upstream of the *TERT* gene, which is oriented towards the telomere (**Fig. 1B**). The breakpoints on the centromere side (BP_cen_) all were located at 16.98 Mb. Both breakpoints fell within TA-rich tandem repeat regions, or hereafter TA-rich sequences, which are difficult to resolve by short-read sequencing due to reduced coverage (**Fig. 1C**) caused by secondary DNA structures and decreased mappability of sequencing reads. The BP_tel_ region spans approximately 700 base pairs (bps) and contains 246 TA dinucleotides, while the BP_cen_ region spans about 130 bps and contains 59 TA dinucleotides. Both regions thus exhibit a density of TA dinucleotides exceeding 35 per 100 bps, substantially higher than the genome-wide average of 6.1 per 100 bps (**Fig. 1C**).

**Figure 1.**
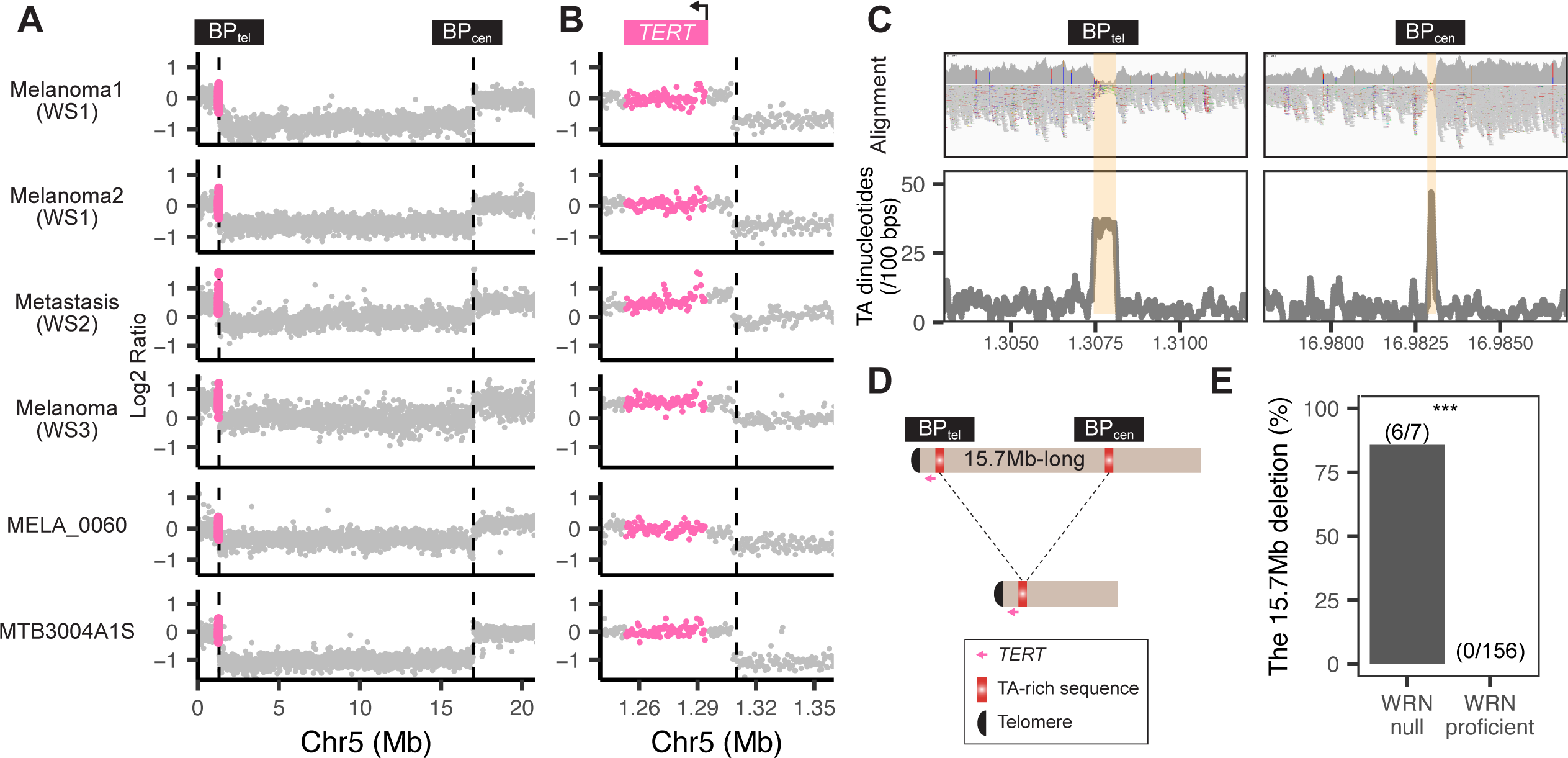
A recurrent deletion on chromosome 5p in melanomas with bi-allelic inactivation of WRN. **(A)** The CNVkit bin-level copy number profiles of chromosome 5p show congruent breakpoints (vertical lines) across six melanomas upstream of *TERT* (highlighted in pink). BP_tel/cen_: breakpoint on the telomeric/centromeric side. **(B)** Close-up view of BP_tel_ and the *TERT* gene. **(C)** Breakpoint regions have an increased density of TA dinucleotides. Upper panel: IGV screenshot with sequencing coverage and mapped reads flanking the breakpoints in melanoma 2 of WS1 as an example. Lower panel: TA dinucleotide density plot across the region shown in the upper panel. **(D)** Diagram showing that the 15.7 Mb-long copy loss was caused by a single deletion rather than two independent structural alterations. **(E)** The deletion is found exclusively in acral and mucosal melanomas with bi-allelic inactivation of WRN (Fisher’s Exact Test, *P* = 2.95 x 10^-10^).

For WS2, we also sequenced five metastases that originated from the same primary tumor, while for WS3, we separately sequenced the *in situ* and invasive areas from the primary tumor as well as a local metastasis. All samples harbored the 15.7 Mb DNA copy loss (**Fig. S2** and **S3**), indicating that the structural rearrangements generating the alterations originated early during the evolution of both tumors.

By examining WGS data of 158 sporadic acral and mucosal melanomas from previous studies (*17*, *18*), we identified two acral melanomas, MELA_0060 and MELA_0309, with biallelic inactivation of the *WRN* gene (**Fig. S4**). In MELA_0060 *WRN* was disrupted in the tumor by the combined effect of a germline deletion of exons 19 to 23 in one allele and somatic loss of the other allele (**Fig. S4**). In MELA_0309, both parental alleles were deleted in the tumor but not germline. Of these two melanomas, MELA_0060 showed an identical 15.7 Mb-long DNA copy loss upstream of *TERT* as the melanomas from the WS patients (**Fig. 1A** and **1B**). The longer insert size of the sequencing library enabled identification of paired-end reads supporting direct joining of BP_tel_ and BP_cen_ in this sample (**Fig. 1D** and **S4**).

From our own institution, we identified another WRN-null sinonasal mucosal melanoma with a truncating germline *WRN* mutation (E388fs) and somatic loss of the remaining allele and established a cell line from it, MTB3004A1S. Short-read and PacBio HiFi long-read WGS of this line revealed loss of the same 15.7 Mb-long region of DNA upstream of *TERT* (**Figs. 1A, 1B** and **S5**). Long reads spanning the TA-rich sequences flanking the deleted region directly captured the junction between BP_tel_ and BP_cen_, confirming that the recurrent 15.7-Mb loss represents a simple deletion (**Fig. 1D** and **S5**).

In summary, an identical 15.7 Mb-long deletion was present in 6 out of 7 separate WRN-null acral/mucosal melanomas, but not in a single case of 156 acral/mucosal melanomas in which *WRN* was intact (**Fig. 1E**; Fisher’s Exact Test, *P* = 2.95 x 10^-10^). This striking association indicates that WRN loss facilitates the formation of this deletion, which must convey a strong selective advantage. Notably, all six WRN-null melanomas with the deletion (**Figs. 1** and **S6**) had at least one inactivated *WRN* allele already in the germline, whereas the only WRN-null tumor without the deletion had somatic inactivation of both *WRN* alleles. This pattern suggests that WRN inactivation promotes genetic alterations that act early during tumor evolution.

### The 15.7 Mb-long deletion causes *TERT* transcriptional activation through enhancer hijacking

We next examined the possible functional consequence of the 15.7 Mb-long deletion upstream of the *TERT* gene. *TERT* is the earliest and most commonly altered gene in acral and mucosal melanomas, activated by structural rearrangements and amplifications (*17–20*). Since the BP_tel_ end of the deletion lies only approximately 12 kb upstream of *TERT*, we reasoned that the deletion may transcriptionally activate *TERT* by changing the genomic and/or chromatin landscape at this locus. We performed ATAC-seq and CUT&Tag profiling of H3K27ac and H3K4me1 in the WRN-null melanoma cell line MTB3004A1S and identified multiple peaks within a 42 kb-long region to the centromeric side of BP_cen_ (**Fig. 2A** and **Data S3**). H3K27ac ChIP-seq datasets from publicly available human melanoma cell lines and melanocytes showed concordant peaks indicating a cluster of enhancers active in the melanocytic lineage (**Fig. S7**). These findings suggested that the 15.7 Mb deletion upstream of the *TERT* gene repositions these enhancers and induces its transcription.

**Figure 2.**
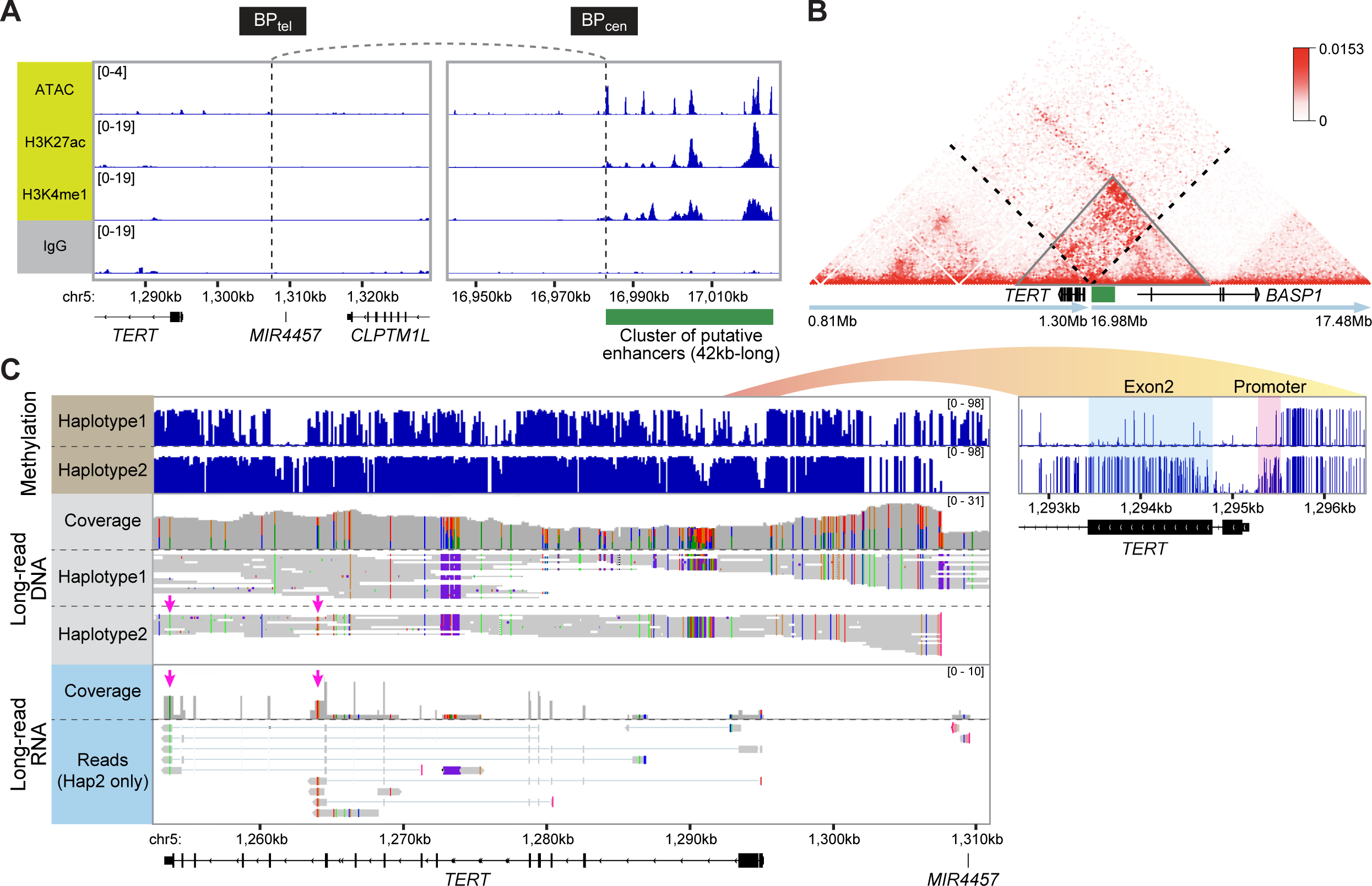
The deletion causes *TERT* transcriptional activation in *cis*, likely through hijacking an upstream enhancer. **A)** ATAC-seq and CUT&Tag signals of H3K27ac, H3K4me1 and IgG control indicate a cluster of active enhancers (green box) downstream of BP_cen_ in the WRN-null cell line MTB3004A1S. **B)** Hi-C contact matrix (5 kb resolution) from MTB3004A1S with dashed black lines indicating the breakpoints flanking the deletion. The green box highlights the 42kb-long region enriched for putative enhancers shown in A. The Hi-C data show increased chromatin contact frequency between this enhancer region and the *TERT* promoter residing within the same TAD (gray triangle). **C)** The haplotype carrying the deletion (haplotype 2) shows allele-specific hypermethylation at the *TERT* promoter and exon 2. Long-read RNA reads map exclusively to haplotype 2 carrying the 15.7 Mb-long deletion as indicated by the SNPs highlighted by pink arrows.

To determine whether *TERT* and the enhancer cluster reside within the same topologically associating domain (TAD) in samples with the deletion, we performed Hi-C sequencing of MTB3004A1S. We called structural variations and mapped TADs onto the rearranged region on chromosome 5p, spanning the deletion. The result demonstrated that *TERT* and the cluster of enhancers reside within a newly formed TAD (neo-TAD) (gray triangle in **Fig. 2B**), supporting a model in which the deletion promotes physical contact of this enhancer region and the *TERT* promoter.

Hypermethylation of the *TERT* promoter, particularly of the hypermethylated oncological region (THOR) upstream of the core promoter, is frequently associated with *TERT* activation in cancers (*21*, *22*). Using haplotype-specific CpG methylation from PacBio HiFi long-read data of MTB3004A1S we found selective hypermethylation across the distal *TERT* promoter exclusively on the haplotype harboring the deletion (haplotype 2, **Fig. 2C**) with additional but less extensive hypermethylation within exon 2 although not exon 1. The first two exons of *TERT* encompass CTCF binding sites, and DNA methylation at these loci can disrupt CTCF occupancy and thereby facilitate promoter activation (*23*, *24*). The haplotype-specific methylation pattern thus strongly supports transcriptional activation of the *TERT* allele in *cis* with deletion.

To test this, we measured allelic-specific expression of *TERT* with Kinnex long-read RNA-seq of MTB3004A1S. All 9 *TERT* transcripts detected were exclusively derived from haplotype 2 (**Fig. 2C**; Binomial test, *P* = 0.0039), supporting monoallelic *TERT* expression from the rearranged haplotype. We further investigated the effect of the rearrangement on *TERT* expression in the melanocytic lineage using AlphaGenome (*25*). We provided either the 1 Mb reference sequence surrounding BP_tel_ (control sequence without the deletion) or a 1 Mb synthetic sequence consisting of equal halves derived from the telomeric side of BP_tel_ and the centromeric side of BP_cen_, representing the expected sequence following the deletion. AlphaGenome predicted increased *TERT* expression in the synthetic sequence compared to predicted expression of the control sequence **(Fig. S8A** and **S8B)**. We further simulated the functional requirement of these elements in silico by masking the entire 42 kb region harboring the candidate enhancer peaks (**Fig. 2A**) with ‘N’ sequences. This large-scale sequence disruption of the rearranged sequence reduced predicted *TERT* expression by more than 50% **(Fig. S8)**. Targeted masking of narrower intervals encompassing only a subset of these peaks similarly compromised *TERT* induction, though the magnitude of the reduction was less pronounced **(Fig. S8)**. These results indicate that the repositioned enhancer cluster at ∼17 Mb drives *TERT* expression in *cis*. Enhancer hijacking was previously proposed as a mechanism of *TERT* activation in these melanoma subtypes on the basis of structural data (*18*, *26*, *27*).

### WRN-null melanomas show an increased incidence of TA-rich sequence-associated rearrangements, disrupting additional driver genes beyond *TERT*

The WRN helicase stabilizes TA-rich regions during DNA replication (*9*). Microsatellite instability (MSI) leads to expansion of TA-repeats, rendering these regions vulnerable to DNA breaks in the context of WRN deficiency (*9*). None of the WRN-null melanomas in our cohort showed evidence of MSI (**Table S1**). The breakpoints of the 15.7 Mb-long deletion upstream of *TERT* were thus located within native, unexpanded TA-rich sequences, indicating that WRN deficiency may also promote breakage at TA-rich sequences in the absence of microsatellite instability or TA-repeat expansion. To investigate this, we assessed the frequency of structural rearrangements within TA-rich sequences throughout the genomes of WRN-deficient tumors.

In whole genome sequencing data WRN-null melanomas harbored on average 12 (**Table S2;** range: 8 - 16) somatic structural rearrangements falling in TA-rich sequences per tumor genome, whereas WRN-proficient melanomas exhibited only 1 (range: 0 - 6) breakpoint (**Fig. 3A**; Wilcoxon rank-sum test, P = 8.9 × 10^-6^). This difference remained significant after accounting for the total number of breakpoints per tumor genome (5.5% vs. 0.1%; **Fig. 3B**; *P* = 1.2 × 10^-5^). Examining the long-read sequencing data of MTB3004A1S for structural rearrangements involving TA-rich sequences, we identified three types of rearrangements: 7 with one end at a TA-rich sequence, 7 with both ends at TA-rich sequences, and 1 in which a TA-rich end was capped by telomeric sequence — 15 events in total (see **Fig. 3E** and **Table S3**). Most of the resulting rearrangements were unbalanced (n=14), with only one balanced rearrangement.

**Figure 3.**
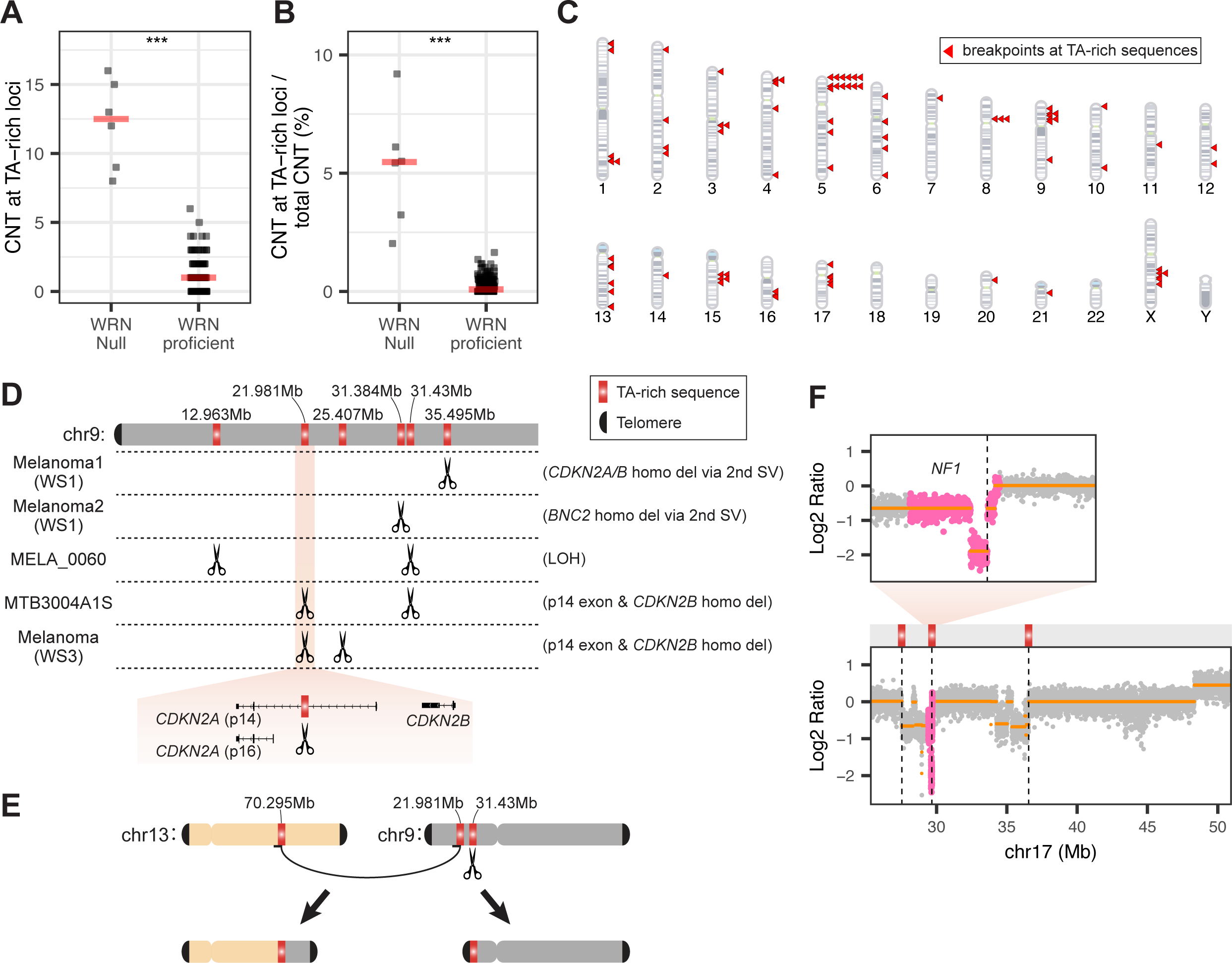
Genome-wide increase of structural rearrangements at TA-rich sequences disrupt melanoma driver genes beyond *TERT*. **A** and **B)** WRN-null melanomas show higher absolute (**A;** Wilcoxon rank-sum test, *P* = 8.9 × 10^-6^) and relative numbers (**B;** *P* = 1.2 × 10^-5^) of somatic copy-number transitions (CNTs) at TA repeats compared to WRN-proficient melanomas. MTB3004A1S was not included considering the lack of a matching germline sample. **C)** The genomic distribution of TA-rich sequence-associated breakpoints across the 7 WRN-null melanomas. Breakpoints observed in more than 1 melanomas are stacked. **D)** Chromosome 9p harbors recurrent breakpoints at six TA-rich loci across melanomas. The upper schematic depicts chromosome 9p and the positions of the six TA repeats. The middle panel shows breakpoint locations (scissors) in individual tumors, with the resulting genomic consequences annotated on the right. MTB3004A1S shares a breakpoint at 21.981 Mb with the WS3 melanoma and a breakpoint at 31.43 Mb with MELA_0060. The bottom panel provides a zoomed-in view of the breakpoint at 21.981 Mb. **E)** The TA-rich sequence at 21.981 Mb on chromosome 9p forms junction with 70.295 Mb on chromosome 13, which exhibited excessive TA-repeats based on long-read data, while the TA-rich sequence at 31.43 Mb on chromosome 9p was capped with telomeric sequences. **F)** Rearrangements result in *NF1* homozygous deletion in melanoma 2 of WS1. The lower panel shows a cluster of breakpoints on chromosome 17q, with three located at TA-rich sequences (vertical dashed lines), while the upper panel shows a zoomed-in view of *NF1*.

Across the seven WRN-null melanomas (**Fig. 1**), a total of 74 TA-rich loci were involved in structural rearrangements, distributed non-randomly across the genome. Besides the two breakpoints flanking the 15.7 Mb-long deletion upstream of *TERT*, nine other TA-rich loci were rearranged in more than one tumor (**Figs. 3C**). The most recurrent cluster affected the *CDKN2A/B* locus on chromosome 9p, where breakpoints at six TA-rich loci recurred across five of the seven tumors (**Fig. 3D**). In these tumors, rearrangement of one allele at a TA-rich sequence, together with loss of the second allele through an independent event not involving a TA-rich sequence, resulted in homozygous deletion of *CDKN2A/B*. MTB3004A1S illustrates the complexity of these events. Its two flanking 9p breakpoints, at 21.981 Mb and 31.43 Mb, both fell within TA-rich sequences but were not joined to each other. The breakpoint at 21.981 Mb, shared with the WS3 melanoma and lying within *CDKN2A*, was fused to a TA-rich sequence at 70.295 Mb on chromosome 13. In contrast, the breakpoint at 31.43 Mb, shared with MELA_0060, was capped by telomeric sequence **(Fig. 3E)**. A 9.45 Mb-long segment spanning *CDKN2A/B* was thereby lost through a complex rearrangement rather than a simple deletion, removing the p14^ARF^-specific exon and *CDKN2B*. In the WS3 melanoma the same 21.981 Mb breakpoint was present, but short-read data could not resolve its partner.

Melanoma 2 of WS1 showed a cluster of breakpoints on chromosome 17q. Three of the breakpoints fell into TA repeats, including one resulting in homozygous deletion of *NF1* with additional rearrangements (**Fig. 3F**).

Together, these data indicate that WRN deficiency increases structural rearrangements at TA-rich sequences genome-wide, with a disproportionate fraction of rearrangements disrupting melanoma-relevant tumor suppressor genes including *CDKN2A, CDKN2B* and *NF1*.

### WRN loss is associated with structural variations at TA-rich sequences across human cancers

To determine whether WRN loss is associated with breakpoints at TA-rich sequences in cancers beyond melanoma, we surveyed WGS data from TCGA. We identified 91 tumors harboring truncating or splice-site *WRN* mutations; these were distributed throughout the coding region, with a recurrent somatic frameshift (S1128fs) and a recurrent germline nonsense mutation (R1406*) (**Fig. 4A**). After excluding samples with low tumor purity or insufficient data quality, 82 tumors were retained (**Data S4**), and partitioned into three groups based on their *WRN* allelic status. Ten tumors, spanning eight cancer types, showed biallelic inactivation — an inactivating germline (n = 4) or somatic (n = 6) alteration on one allele together with loss of the second allele in the tumor — and are hereafter referred to as WRN-null. A further 70 tumors carried a single truncating or splice-site mutation with no detectable second hit (monoallelic). The remaining 2 tumors each harbored two truncating mutations that could not be phased as *cis* or *trans*, leaving their allelic status unresolved; these were excluded from downstream analyses. None of the 10 WRN-null tumors were melanomas and none carried the recurrent 15.7 Mb-long deletion upstream of *TERT*.

**Figure 4.**
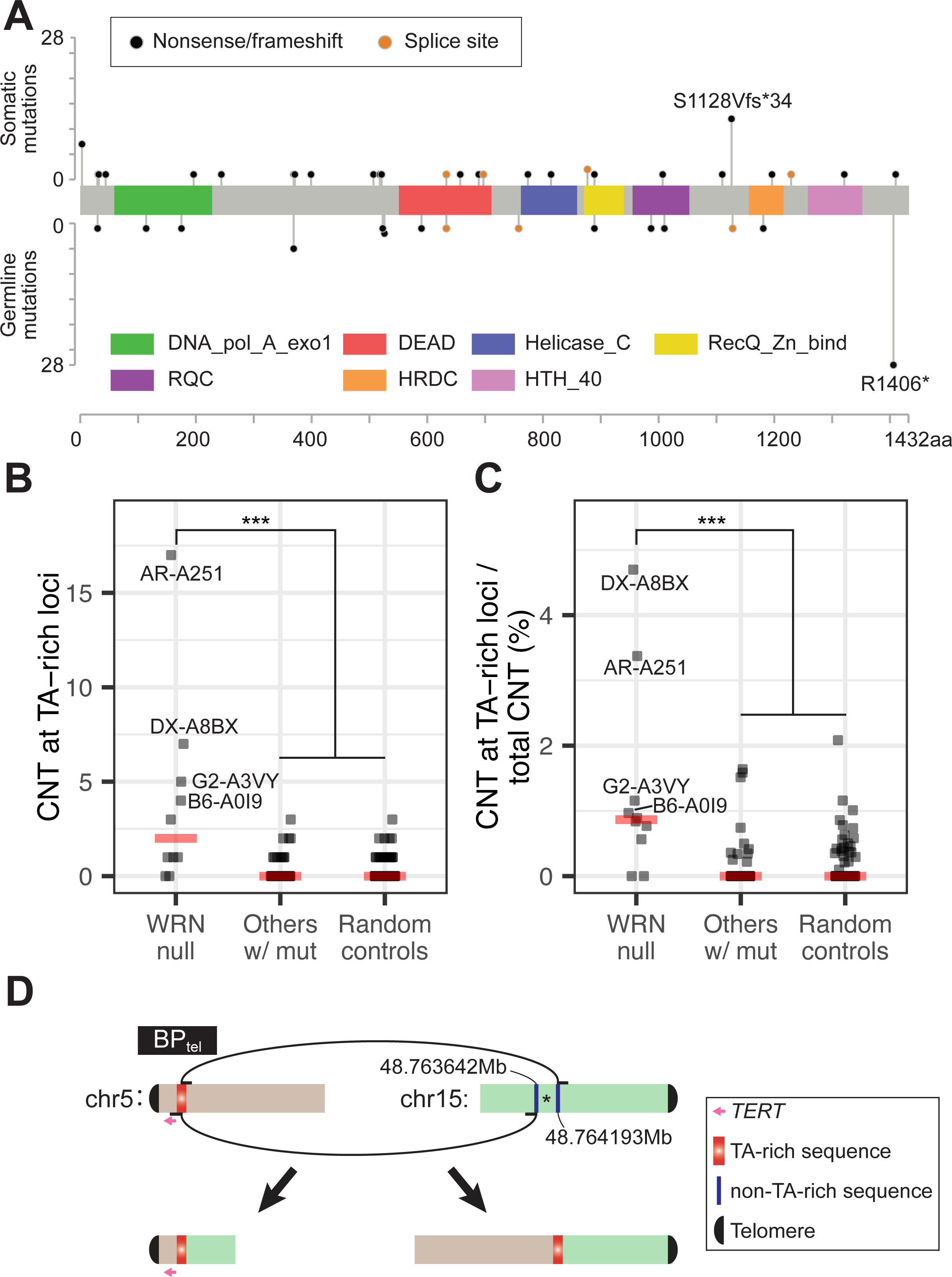
WRN loss is associated with rearrangements at TA-rich sequences in non-melanoma cancers. **A)** Lollipop plot showing the landscape of truncating *WRN* mutations in cancers from TCGA. B and C) Tumors with biallelic WRN inactivation exhibit higher absolute (B) and breakpoint-normalized (C) numbers of somatic copy-number transitions (CNTs) involving TA-rich sequences than tumors harboring WRN truncating mutations without evidence of biallelic inactivation or randomly selected control tumors (Wilcoxon rank-sum test; B, *P* = 1.63 × 10^-6^ and 1.0 × 10^-4^, respectively; C, *P* = 1.15 × 10^-6^ and 3.91 × 10^-5^, respectively). D) Rearrangements involving BP_tel_ in DX-A8BX. The segment marked with an asterisk (*) on chromosome 15 was involved in a separate rearrangement not shown here. Genomic coordinates are based on hg19.

The group of tumors with bi-allelic loss of WRN harbored significantly more rearrangements at TA-rich sequences than the group with monoallelic alterations (**Fig. 4B** and **Table S4**; median 2 vs. 0; *P* = 1.63 × 10^-6^) or a group of 80 randomly selected tumors from the same eight cancer types (10 per type; median 0; *P* = 1.0 × 10^-4^), and this difference remained after normalizing for each tumor’s total rearrangement burden (**Fig. 4C**; median 0.87% vs. 0; P = 1.15 × 10^-6^ and 3.91 × 10^-5^, respectively). As in melanoma, rearrangements involving TA-rich sequences were most frequent in tumors with germline *WRN* inactivation: the four tumors with the highest burden (≥4 breakpoints at TA-rich sequences) — AR-A251 and B6-A0I9 (breast invasive ductal carcinoma), DX-A8BX (undifferentiated pleomorphic sarcoma from the retroperitoneum), and G2-A3VY (bladder urothelial carcinoma) — each carried a germline first hit. No breakpoint was shared across this smaller cohort. Notably, the sarcoma harbored a copy-number-balanced rearrangement at the BP_tel_ locus 12 kb upstream of *TERT* identified in the WRN-null melanomas. However, the partner of this rearrangement was a region of chromosome 15 predicted to act as an enhancer in the mesenchymal lineage (**Fig. 4D** and **Fig. S9**). Thus, the same fragile site upstream of *TERT* was engaged in a second, unrelated WRN-deficient tumor type, where it also juxtaposed *TERT* to an active enhancer. This suggests that WRN loss generates rearrangements at TA-rich sequences beyond melanoma, and that this breakpoint upstream of *TERT* may be recurrently engaged in more than one lineage.

## Discussion

Our findings redefine the role of WRN in cancer. WRN is not merely a general guardian of genome stability whose loss broadly increases instability; it is a specific suppressor of structural rearrangement at TA-rich tandem repeats. When WRN is lost, these sequences become prone to rearrangement, which can restructure the genome and disrupt cancer genes. This differs fundamentally from the established role of WRN at repetitive sequence. In mismatch-repair-deficient cancers, tandem repeats become WRN-dependent only after they are somatically expanded in length. By contrast, the tumors we describe are microsatellite-stable with no evident change in tract lengths. While the lengths of TA-rich sequences vary in the population, the near-complete association of the 15.7 Mb-long deletion upstream of *TERT* with WRN loss argues against a requirement for a specific constitutional configuration of the repeats. Otherwise, their rearrangement would be unlikely to track so closely with WRN status.

The detailed process generating these rearrangements is unresolved. In MTB3004A1S, 7 of 15 rearrangements involving TA-rich sequences joined to a second TA-rich sequence, far exceeding the ∼0.2% of the genome these repeats occupy and indicating that TA-rich sequences are strongly favored as rearrangement partners. How these junctions form remains open: they are compatible with the joining of concomitant double-strand breaks or with replication-based template switching, both consistent with WRN’s function at replication forks.

The narrow tumor spectrum in WS likely follows from this mutational mechanism. WRN loss generates rearrangements at TA-rich sequences genome-wide, but the tumors that arise are those in which such a rearrangement alters a gene whose deregulation confers a selective advantage early in a given tissue. Acral and mucosal melanoma illustrate this coincidence. In these subtypes, *TERT* activation is the earliest and most prevalent driver event, whereas cutaneous melanomas are initiated by *BRAF* or *NRAS* mutation, with *TERT* activated only later (*28*, *29*). The near-exclusive occurrence of the 15.7 Mb-long deletion upstream of *TERT* in tumors that harbor germline *WRN* loss reflects this: only when WRN is absent from the outset can the deletion provide an early, rate-limiting oncogenic alteration in these melanomas. In cutaneous melanoma by contrast, *TERT* activation is not an early requirement, and WRN deficiency accordingly does not confer susceptibility. The specific 15.7 Mb-long deletion was melanoma-specific, likely because the enhancer it captures is active in melanocytes. But the underlying route — BP_tel_ fragility followed by capture of a lineage-active enhancer — may be more general: in the WRN-deficient sarcoma, the same BP_tel_ breakpoint instead joined chromosome 15q, adjacent to a region predicted to act as a mesenchymal enhancer, in a configuration that may similarly engage *TERT*.

Given WRN’s role in resolving non-B DNA structures during replication, the TA-rich sequences resolved here are likely one instance of a wider susceptibility, extending to other structure-prone sequences that depend on WRN.

## Materials and Methods

### Specimens from WS patients

For the three WS patients (WS1, WS2 and WS3), we collected archival formalin-fixed and paraffin-embedded (FFPE) tissue specimens. WS1 and WS2 were a pair of Japanese sisters (*14*, *15*). For WS1, two primary mucosal melanomas, each from the left and right nasal cavity respectively, were obtained. Notably, this patient was also diagnosed with another three primary melanoma lesions (one from pudendum and two from left plantar), as well as a fourth primary melanoma from the right sole (*14*); however, tissue specimens from these lesions were not available for analysis. For WS2, the primary tumor was located in the left nasal cavity. This primary tumor was excluded from analysis due to prior decalcification. Instead, we included five metastatic lesions (from the abdomen, liver, neck, right forearm, and waist) of the same melanoma. WS3 was from a different patient (*16*). We collected two primary tumor areas, a melanoma *in situ* and an invasive, from right heel, as well as a local metastasis from the right medial ankle. In total, ten tumor samples representing four different melanomas and three adjacent normal samples were included from the three patients. All human specimens were studied under institutional review board approval, and the contributing sites’ ethics approvals and participant consents are in place for the analyses reported here.

### DNA extraction and short-read whole-genome sequencing

Five-µm hematoxylin and eosin-stained (H&E) sections of FFPE tissues were reviewed by two pathologists to identify tumor and adjacent non-lesional areas. Microdissection of these areas was performed on 10 µm unstained sections under a dissection microscope (SMZ-171-BLED, Motic). The tissue sections were processed for dual DNA and RNA extractions using the CELLDATA DNAstorm/RNAstorm 1.0 Combination Kit (CD508, Biotium) and quantified using the Qubit 2.0 Fluorometer with the dsDNA High Sensitivity assay Kit (Q32854, Thermo Fisher Scientific). 250 - 330 ng of genomic DNA was sheared to a median fragment length of 200 bp using a Covaris E220 Sonicator. The fragmented double-stranded DNA was repaired using the NEBNext FFPE DNA Repair Mix v1 (M6630L, NEB). DNA libraries were prepared using the KAPA HyperPrep Kit (KK8505, Roche) and quantified in a LightCycler 480 II System (Roche) by quantitative PCR (qPCR) using the KAPA Library Quantification Kit (KK4854, Roche). The size distribution was determined using an Agilent 2100 Bioanalyzer with the High Sensitivity DNA Kit (5067–4626, Agilent). The libraries were then sequenced on an Illumina NovaSeq X with 150 bp paired-end reads.

### Processing of WGS sequencing data

After obtaining the FASTQ sequencing data, we examined the sequencing quality using FastQC and trimmed 3’ adaptors using fastp (v0.23.4) (*30*). The trimmed reads were then aligned to the hg19 genome assembly using the Burrows-Wheeler Aligner (v0.7.17) (*31*) to generate BAM files. Picard/GATK were used to remove PCR duplicates, calculate insert size and sequencing coverage, perform base quality score recalibration and realign indel regions. We used Mutect2 (*9*) to call somatic point mutations and indels and ANNOVAR (*32*) for annotation. The mutations were filtered by requiring: 1) >= 4 reads supporting the mutated allele; 2) the mutant allele frequency is >= 5%; 3) the sequencing depth of both tumor and normal samples being >= 10; 4) the population frequency of the mutation being <= 1% in any of the 5 databases (ExAC, GnomAD, 1000 Genome, dbSNP150, and NHLBI-ESP). Germline mutations were called using HaplotypeCaller against tumor adjacent samples and annotated using ANNOVAR. Copy number analyses were performed using CNVkit (v0.9.12) (*33*) and FACETS (v0.6.2) (*34*). Microsatellite instability (MSI) score was estimated using MSIsensor2 (v0.1) (*35*).

### Short-read WGS of the MTB3004A1S cell line

The MTB3004A1S cell line was established from a sinonasal mucosal melanoma in our recent study (*36*). We extracted DNA from 2 million cells using the Qiagen DNeasy Blood & Tissue Kit (69504, Qiagen). Quantification, quality control, sequencing, alignment, and BAM file processing were performed as described above for FFPE samples. Mutations were called using Mutect2 tumor-only mode and copy number analyses were performed using CNVkit with a flat reference.

### Pacbio long-read WGS and analysis of MTB3004A1S cell line

High molecular weight (HMW) DNA was extracted from a pellet of 2.5 million cells using the Nanobind CBB Kit (102-301-900, PacBio). The DNA was quantified using the Qubit 2.0 Fluorometer with the Qubit dsDNA BR Assay Kit (Q32850, Thermo Fisher Scientific). We sheared 2 mg of DNA into approx. 15 kb fragments using centrifugal force by applying the sample to a Covaris g-TUBE column (520079, Covaris) and running in an Eppendorf 5424 Centrifuge at 3200 rpm, four times, for 16 min per pass. Library preparation was performed using the SMRTbell Prep Kit 3.0 (102-178-400, Pacific Biosciences). Fragment length distributions of the sheared DNA and the DNA library were evaluated using the FemtoPulse System (M5330AA, Agilent Technologies) with the gDNA 165kb analysis Kit (FP-1002-0275, Agilent Technologies). The library was then sequenced on a single SMRT cell using the PacBio Revio system.

After obtaining HiFi long reads, we aligned onto the hg19 genome reference using pbmm2 (v1.13.1). We then used DeepVariant (v1.8.0) to call variants (*37*), followed by WhatsHap (v2.3) to phase haplotypes (*38*). DNA methylation levels for each haplotype were then estimated with pb-CpG-tools (v2.3.2). Structural variants were identified using pbsv (v2.9.0), and copy-number profiles were additionally assessed with Delly CNV (*39*).

### Pacbio long-read RNA-seq and analysis of MTB3004A1S cell line

RNA library preparation was performed by the Center for Advanced Technology (CAT) core at UCSF using the Kinnex Full-Length RNA Kit (PacBio) according to the manufacturer’s protocol, with MTB3004A1S and 3 other cell lines multiplexed into the same library. The library was sequenced on a SMRT cell of the PacBio Revio system. After generating HiFi reads, we first applied Skera (v1.4.0) to segment MAS-Seq concatenated reads using the concatenation adapters. We then performed sample demultiplexing with lima (v2.13.0), followed by isoseq (v4.3.0) to remove poly(A) tails and synthetic concatemers. The resulting full-length reads were aligned to the hg19 reference using pbmm2.

### Cut&Tag, ATAC-seq and analysis of MTB3004A1S

CUT&Tag of H3K27ac, H3K4me1 and IgG, and ATAC-seq library preparation and sequencing were performed by Active Motif (Carlsbad, CA) according to the manufacturer’s standard protocols.

For both CUT&Tag and ATAC-seq data, we used fastp to trim 3’ adaptor, followed by bowtie2 to map onto the hg19 reference genome. We then sorted the mapped reads, and marked duplicates. For ATAC-seq, we further removed reads mapped onto mitochondria or regions from ENCODE blacklist, and shifted the mapped reads used alignmentSieve from deepTools (--ATACshift) (v3.5.6) (*40*). We then used bamCoverage to generate bigwig files, with the CPM normalization method. Peaks were called using MACS3 (v3.0.2) hmmratac for ATAC-seq sample (*41*), and SEACR (v1.3) for Cut&Tag data with IgG sample as control (*42*).

### Analyses of public ChIP-seq data

We obtained the raw H3K27ac ChIP-seq data of melanoma cell lines WM35, WM1552, YUPEET and YUCHIME from Mendelson et al (*43*) and SK-MEL-5 from Takahashi et al (*44*). Input ChIP-seq data were available for all except SK-MEL-5. We processed all datasets using the same pipeline described above for the CUT&Tag data, including alignment, sorting, deduplication, and bigWig file generation, independently for both H3K27ac and input samples. Peaks were then called using MACS3.

### Hi-C and analysis

Hi-C library preparation and sequencing were performed by Arima Genomics (Carlsbad, CA) according to the manufacturer’s standard protocols. We aligned sequencing reads to the hg19 reference genome using BWA (bwa mem-SP5M). Pairtools (v1.1.2) (*45*) was then used to parse the raw BAM files with the walking policy --5unique, sort alignments, remove duplicates, and retain read pairs of types “UU”, “UR”, “RU”, or “uu”. Cooler was subsequently used to generate the .mcool file. To identify neo-TADs, we used newtad-caller implemented in NeoLoopFinder (v0.4.3-r2) (*46*), following prior steps of CNV estimation, SV detection, and complex SV assembly. SVs predicted at 5 kb resolution using EagleC (v0.1.9) (*47*) were incorporated.

### Application of AlphaGenome

To evaluate the effect of the 15.7 Mb-long deletion in the melanocytic lineage, we used foreskin melanocytes (Cell Ontology term: CL:2000045), the only cell type in AlphaGenome with both RNA-seq and multiple layers of chromatin profiling data available. We provided either a 1 Mb reference sequence surrounding BPtel (representing the control sequence without the deletion) or a 1 Mb synthetic sequence consisting of equal halves derived from the telomeric side of BPtel and the centromeric side of BPcen (representing the expected sequence following the deletion). We further simulated the functional requirement of these regulatory elements in silico by masking the entire 42-kb region harboring the candidate enhancer peaks with N sequences, as well as narrower intervals encompassing subsets of these peaks. We additionally applied AlphaGenome to predict the H3K27ac profile surrounding the two breakpoints (BP_tel_ and its partner breakpoint on chromosome 15q) of the translocation observed in the undifferentiated pleomorphic sarcoma DX-A8BX, using available mesenchymal lineage cell types, including mesenchymal stem cells (CL:0000134), dermal fibroblasts (CL:0002551), lung fibroblasts (CL:0002557), and osteoblast (CL:0000062).

### Identifying breakpoints located at TA-rich sequences through WGS data

To first annotate TA-rich sequences in the human reference genome, we performed a sliding-window scan using a 50-bp window and a 5-bp step size for each chromosome. For windows with zero ambiguous (“N”) bases and at least 40 A/T nucleotides, we further filtered to retain those satisfying at least one of the following conditions: 1) containing ≥1 tract of 10 consecutive TA dinucleotides; 2) containing ≥1 tract of 6 consecutive TA dinucleotides and a ≥40-bp palindromic segment within the same window; 3) containing ≥15 TA dinucleotides in total. After identifying all qualifying windows, we merged neighboring windows located within 500 bp of each other into larger candidate regions. Finally, we removed regions with predicted folding free energy (calculated using NUPACK v4.0.1.12 (*48*)) greater than −10 kcal/mol. In total, 31502 such regions were identified.

To identify copy-number breakpoints occurring at TA-rich sequences, we applied several filtering steps to remove false positives. First, we obtained bin-level log₂ ratio values from CNVkit using 500 bp bins. For each TA-rich sequence locus, we compared the log₂ ratio difference between bins within the 10 kb upstream and 10 kb downstream flanking regions using Wilcoxon rank sum test. An FDR threshold of < 0.05 was then applied. Next, we removed TA-rich sequences located more than 1 Kb away from all FACETS-detected copy-number breakpoints. For the remaining candidates, we manually inspected each site and excluded TA-rich sequences: i) with a clear breakpoint located very close to, but outside of, the repeat (e.g., within tens to hundreds of base pairs), and ii) with evidence of mapping artifacts in the flanking regions, such as excessive reads with mapping quality 0 or high mismatch rates. The remaining TA-rich sequences were retained as bona fide sites containing copy-number breakpoints.

For MTB3004A1S, we examined SVs identified from long-read data and required that at least one breakpoint be located within a TA-rich sequence. To account for potential inter-individual variability in TA-rich sequence length, and because some TA-rich sequence tracts can extend for several kilobases, non-translocation alterations (e.g., inversions and deletions) were required to span at least 10 kb. We further manually reviewed the two breakpoints of each junction through IGV to ensure that they were not located within the same TA-rich sequence. To further identify breakpoints that may be missed by pbsv, we separately examined the bin-level log2 ratios from CNVkit in the same manner as described above, and also manually reviewed TA-rich sequences that harbored breakpoints in the other WRN-null melanomas.

### Examining WGS samples from TCGA

To identify tumors harboring germline or somatic truncating or canonical splice-site mutations in *WRN*, we used the BAM-slicing function of the Genomic Data Commons (GDC) Data Portal (*49*) to retrieve sequencing reads spanning the *WRN* locus from all TCGA WGS samples. Variants were called using GATK HaplotypeCaller and annotated with ANNOVAR. For samples carrying at least one truncating or canonical splice-site mutation, we downloaded the matched tumor and normal WGS data and performed copy-number analysis using CNVkit and FACETS to determine whether *WRN* had undergone biallelic inactivation in the tumor. This analysis identified 10 tumors, representing eight cancer types, with biallelic *WRN* inactivation. For each cancer type, we additionally selected 10 tumor-normal pairs at random as controls. TA-rich sequence-associated structural breakpoints were then identified using the same pipeline described above and compared among WRN-null tumors, tumors with monoallelic WRN inactivation, and the random control cohort. To match the hg38 reference used for the TCGA data, we converted the hg19 coordinates of the TA-rich sequence regions identified above to hg38 using the UCSC LiftOver tool. Unless otherwise noted, all presented data were based on the hg19 reference.

## Acknowledgments

We thank all the included patients from which we obtained specimens and sequencing data. We also thank Satoshi Fukushima for facilitating sample collection, and members in the Bastian and Yeh labs for helpful discussion. Sequencing was performed at the Center for Advanced Technology (CAT) of UCSF.

## Author contributions

Conceptualization: B.C.B., M.W., M.G., I.Y.

Methodology: M.W., H.P., A.S., A.J.E., N.C.P., M.G., I.Y., B.C.B.

Investigation: M.W., H.P., A.S., A.J.E., N.C.P., S.M., J.D.A., K.K.T., A.I.D., I.E.S., H.N., S.F., T.H., S.P., S.D., A.O., I.Y., B.C.B.

Visualization: M.W.

Writing – original draft: M.W., B.C.B.

Writing – review & editing: M.W., B.C.B., A.H.S., I.Y.

## Data, code, and materials availability

Processed data supporting the findings, including somatic mutations, copy-number profiles, and breakpoint calls, are provided in the Supplementary Information and will also be deposited in the European Genome-phenome Archive (EGA). Raw whole-genome sequencing reads are available under controlled access through EGA, with the exception of WS3 and MTB3004A1S, for which participant consent and institutional approvals restrict the deposition of germline-containing sequence data. Accordingly, raw sequencing reads for these samples are not deposited, and somatic-level processed data are provided instead. Requests for access to restricted data will be considered in accordance with the governing consent and institutional approvals.

## References

1. M. Goto, R. W. Miller, Y. Ishikawa, H. Sugano, Excess of rare cancers in Werner syndrome (adult progeria). Cancer Epidemiol Biomarkers Prev 5, 239–246 (1996).

2. J. M. Lauper, A. Krause, T. L. Vaughan, R. J. M. Jr, Spectrum and Risk of Neoplasia in Werner Syndrome: A Systematic Review. PLOS ONE 8, e59709 (2013).

3. M. Muftuoglu, J. Oshima, C. von Kobbe, W.-H. Cheng, D. F. Leistritz, V. A. Bohr, The clinical characteristics of Werner syndrome: molecular and biochemical diagnosis. Hum Genet 124, 369–377 (2008).

4. J. Oshima, J. M. Sidorova, R. J. Monnat, Werner syndrome: Clinical features, pathogenesis and potential therapeutic interventions. Ageing Research Reviews 33, 105–114 (2017).

5. C.-E. Yu, J. Oshima, Y.-H. Fu, E. M. Wijsman, F. Hisama, R. Alisch, S. Matthews, J. Nakura, T. Miki, S. Ouais, G. M. Martin, J. Mulligan, G. D. Schellenberg, Positional Cloning of the Werner’s Syndrome Gene. Science 272, 258–262 (1996).

6. R. M. Brosh, J. Waheed, J. A. Sommers, Biochemical Characterization of the DNA Substrate Specificity of Werner Syndrome Helicase*. Journal of Biological Chemistry 277, 23236–23245 (2002).

7. M. Fry, L. A. Loeb, Human Werner Syndrome DNA Helicase Unwinds Tetrahelical Structures of the Fragile X Syndrome Repeat Sequence d(CGG)n*. Journal of Biological Chemistry 274, 12797–12802 (1999).

8. L. Crabbe, R. E. Verdun, C. I. Haggblom, J. Karlseder, Defective Telomere Lagging Strand Synthesis in Cells Lacking WRN Helicase Activity. Science 306, 1951–1953 (2004).

9. N. van Wietmarschen, S. Sridharan, W. J. Nathan, A. Tubbs, E. M. Chan, E. Callen, W. Wu, F. Belinky, V. Tripathi, N. Wong, K. Foster, J. Noorbakhsh, K. Garimella, A. Cruz-Migoni, J. A. Sommers, Y. Huang, A. A. Borah, J. T. Smith, J. Kalfon, N. Kesten, K. Fugger, R. L. Walker, E. Dolzhenko, M. A. Eberle, B. E. Hayward, K. Usdin, C. H. Freudenreich, R. M. Brosh, S. C. West, P. J. McHugh, P. S. Meltzer, A. J. Bass, A. Nussenzweig, Repeat expansions confer WRN dependence in microsatellite-unstable cancers. Nature 586, 292–298 (2020).

10. E. M. Chan, T. Shibue, J. M. McFarland, B. Gaeta, M. Ghandi, N. Dumont, A. Gonzalez, J. S. McPartlan, T. Li, Y. Zhang, J. B. Liu, J.-B. Lazaro, P. Gu, C. G. Piett, A. Apffel, S. O. Ali, R. Deasy, P. Keskula, R. W. S. Ng, E. A. Roberts, E. Reznichenko, L. Leung, M. Alimova, M. Schenone, M. Islam, Y. E. Maruvka, Y. Liu, J. Roper, S. Raghavan, M. Giannakis, Y.-Y. Tseng, Z. D. Nagel, A. D’Andrea, D. E. Root, J. S. Boehm, G. Getz, S. Chang, T. R. Golub, A. Tsherniak, F. Vazquez, A. J. Bass, WRN helicase is a synthetic lethal target in microsatellite unstable cancers. Nature 568, 551–556 (2019).

11. F. M. Behan, F. Iorio, G. Picco, E. Gonçalves, C. M. Beaver, G. Migliardi, R. Santos, Y. Rao, F. Sassi, M. Pinnelli, R. Ansari, S. Harper, D. A. Jackson, R. McRae, R. Pooley, P. Wilkinson, D. van der Meer, D. Dow, C. Buser-Doepner, A. Bertotti, L. Trusolino, E. A. Stronach, J. Saez-Rodriguez, K. Yusa, M. J. Garnett, Prioritization of cancer therapeutic targets using CRISPR–Cas9 screens. Nature 568, 511–516 (2019).

12. L. Kategaya, S. K. Perumal, J. H. Hager, L. D. Belmont, Werner Syndrome Helicase Is Required for the Survival of Cancer Cells with Microsatellite Instability. iScience 13, 488–497 (2019).

13. S. Lieb, S. Blaha-Ostermann, E. Kamper, J. Rippka, C. Schwarz, K. Ehrenhöfer-Wölfer, A. Schlattl, A. Wernitznig, J. J. Lipp, K. Nagasaka, P. van der Lelij, G. Bader, M. Koi, A. Goel, R. A. Neumüller, J.-M. Peters, N. Kraut, M. A. Pearson, M. Petronczki, S. Wöhrle, Werner syndrome helicase is a selective vulnerability of microsatellite instability-high tumor cells. eLife 8, e43333 (2019).

14. H. Shibuya, A. Kato, N. Kai, S. Fujiwara, M. Goto, A Case of Werner Syndrome with Three Primary Lesions of Malignant Melanoma. The Journal of Dermatology 32, 737–744 (2005).

15. Y. Kadowaki, S. Kodama, M. Moriyama, M. Suzuki, The Relationship between Werner Syndrome and Sinonasal Malignant Melanoma: Two Sibling Cases of Werner Syndrome with Malignant Melanoma. Case Reports in Otolaryngology 2017, e9361612 (2017).

16. A. McGrath, M. Lockhart, T. Griffin, S. A. Lynch, S. F. Dinneen, ‘Werner Syndrome foot’—A case series of four Irish Traveller siblings with Werner Syndrome, diabetes mellitus and complex foot disease. Diabetic Medicine 41, e15390 (2024).

17. F. Newell, Y. Kong, J. S. Wilmott, P. A. Johansson, P. M. Ferguson, C. Cui, Z. Li, S. H. Kazakoff, H. Burke, T. J. Dodds, A.-M. Patch, K. Nones, V. Tembe, P. Shang, L. van der Weyden, K. Wong, O. Holmes, S. Lo, C. Leonard, S. Wood, Q. Xu, R. V. Rawson, P. Mukhopadhyay, R. Dummer, M. P. Levesque, G. Jönsson, X. Wang, I. Yeh, H. Wu, N. Joseph, B. C. Bastian, G. V. Long, A. J. Spillane, K. F. Shannon, J. F. Thompson, R. P. M. Saw, D. J. Adams, L. Si, J. V. Pearson, N. K. Hayward, N. Waddell, G. J. Mann, J. Guo, R. A. Scolyer, Whole-genome landscape of mucosal melanoma reveals diverse drivers and therapeutic targets. Nature Communications 10, 3163 (2019).

18. F. Newell, J. S. Wilmott, P. A. Johansson, K. Nones, V. Addala, P. Mukhopadhyay, N. Broit, C. M. Amato, R. Van Gulick, S. H. Kazakoff, A.-M. Patch, L. T. Koufariotis, V. Lakis, C. Leonard, S. Wood, O. Holmes, Q. Xu, K. Lewis, T. Medina, R. Gonzalez, R. P. M. Saw, A. J. Spillane, J. R. Stretch, R. V. Rawson, P. M. Ferguson, T. J. Dodds, J. F. Thompson, G. V. Long, M. P. Levesque, W. A. Robinson, J. V. Pearson, G. J. Mann, R. A. Scolyer, N. Waddell, N. K. Hayward, Whole-genome sequencing of acral melanoma reveals genomic complexity and diversity. Nature Communications 11, 5259 (2020).

19. I. Yeh, E. Jorgenson, L. Shen, M. Xu, J. P. North, A. H. Shain, D. Reuss, H. Wu, W. A. Robinson, A. Olshen, A. von Deimling, P.-Y. Kwok, B. C. Bastian, M. M. Asgari, Targeted Genomic Profiling of Acral Melanoma. J Natl Cancer Inst 111, 1068–1077 (2019).

20. M. Wang, S. Fukushima, Y.-S. Sheen, E. Ramelyte, N. Cruz-Pacheco, C. Shi, S. Liu, I. Banik, J. D. Aquino, M. Sangueza Acosta, M. Levesque, R. Dummer, J.-Y. Liau, C.-Y. Chu, A. H. Shain, I. Yeh, B. C. Bastian, The genetic evolution of acral melanoma. Nat Commun 15, 6146 (2024).

21. X. Yuan, C. Larsson, D. Xu, Mechanisms underlying the activation of TERT transcription and telomerase activity in human cancer: old actors and new players. Oncogene 38, 6172–6183 (2019).

22. D. D. Lee, M. Komosa, N. M. Nunes, U. Tabori, DNA methylation of the *TERT* promoter and its impact on human cancer. Current Opinion in Genetics & Development 60, 17–24 (2020).

23. S. Renaud, D. Loukinov, F. T. Bosman, V. Lobanenkov, J. Benhattar, CTCF binds the proximal exonic region of hTERT and inhibits its transcription. Nucleic Acids Research 33, 6850–6860 (2005).

24. S. Renaud, D. Loukinov, Z. Abdullaev, I. Guilleret, F. T. Bosman, V. Lobanenkov, J. Benhattar, Dual role of DNA methylation inside and outside of CTCF-binding regions in the transcriptional regulation of the telomerase hTERT gene. Nucleic Acids Research 35, 1245–1256 (2007).

25. Ž. Avsec, N. Latysheva, J. Cheng, G. Novati, K. R. Taylor, T. Ward, C. Bycroft, L. Nicolaisen, E. Arvaniti, J. Pan, R. Thomas, V. Dutordoir, M. Perino, S. De, A. Karollus, A. Gayoso, T. Sargeant, A. Mottram, L. H. Wong, P. Drotár, A. Kosiorek, A. Senior, R. Tanburn, T. Applebaum, S. Basu, D. Hassabis, P. Kohli, Advancing regulatory variant effect prediction with AlphaGenome. Nature 649, 1206–1218 (2026).

26. W. S. Liang, W. Hendricks, J. Kiefer, J. Schmidt, S. Sekar, J. Carpten, D. W. Craig, J. Adkins, L. Cuyugan, Z. Manojlovic, R. F. Halperin, A. Helland, S. Nasser, C. Legendre, L. H. Hurley, K. Sivaprakasam, D. B. Johnson, H. Crandall, K. J. Busam, V. Zismann, V. Deluca, J. Lee, A. Sekulic, C. E. Ariyan, J. Sosman, J. Trent, Integrated genomic analyses reveal frequent TERT aberrations in acral melanoma. Genome Res. 27, 524–532 (2017).

27. F. Newell, P. A. Johansson, J. S. Wilmott, K. Nones, V. Lakis, A. L. Pritchard, S. N. Lo, R. V. Rawson, S. H. Kazakoff, A. J. Colebatch, L. T. Koufariotis, P. M. Ferguson, S. Wood, C. Leonard, M. H. Law, K. M. Brooks, N. Broit, J. M. Palmer, K. L. Couts, I. A. Vergara, G. V. Long, A. P. Barbour, O. E. Nieweg, B. Shivalingam, W. A. Robinson, J. R. Stretch, A. J. Spillane, R. P. M. Saw, K. F. Shannon, J. F. Thompson, G. J. Mann, J. V. Pearson, R. A. Scolyer, N. Waddell, N. K. Hayward, Comparative Genomics Provides Etiologic and Biological Insight into Melanoma Subtypes. Cancer Discovery 12, 2856–2879 (2022).

28. A. H. Shain, I. Yeh, I. Kovalyshyn, A. Sriharan, E. Talevich, A. Gagnon, R. Dummer, J. North, L. Pincus, B. Ruben, W. Rickaby, C. D’Arrigo, A. Robson, B. C. Bastian, The Genetic Evolution of Melanoma from Precursor Lesions. New England Journal of Medicine 373, 1926–1936 (2015).

29. A. H. Shain, N. M. Joseph, R. Yu, J. Benhamida, S. Liu, T. Prow, B. Ruben, J. North, L. Pincus, I. Yeh, R. Judson, B. C. Bastian, Genomic and Transcriptomic Analysis Reveals Incremental Disruption of Key Signaling Pathways during Melanoma Evolution. Cancer Cell 34, 45–55.e4 (2018).

30. S. Chen, Y. Zhou, Y. Chen, J. Gu, fastp: an ultra-fast all-in-one FASTQ preprocessor. Bioinformatics 34, i884–i890 (2018).

31. H. Li, R. Durbin, Fast and accurate long-read alignment with Burrows–Wheeler transform. Bioinformatics 26, 589–595 (2010).

32. K. Wang, M. Li, H. Hakonarson, ANNOVAR: functional annotation of genetic variants from high-throughput sequencing data. Nucleic Acids Research 38, e164–e164 (2010).

33. E. Talevich, A. H. Shain, T. Botton, B. C. Bastian, CNVkit: Genome-Wide Copy Number Detection and Visualization from Targeted DNA Sequencing. PLOS Computational Biology 12, e1004873 (2016).

34. R. Shen, V. E. Seshan, FACETS: allele-specific copy number and clonal heterogeneity analysis tool for high-throughput DNA sequencing. Nucleic Acids Res 44, e131–e131 (2016).

35. B. Niu, K. Ye, Q. Zhang, C. Lu, M. Xie, M. D. McLellan, M. C. Wendl, L. Ding, MSIsensor: microsatellite instability detection using paired tumor-normal sequence data. Bioinformatics 30, 1015–1016 (2014).

36. I. Banik, S. Basu, M. Wang, A. J. Escobar, V. Steri, J. A. Camara Serrano, P. Phojanakong, N. C. Pacheco, K. K. Tsai, E. A. Alvarez, A. Daud, I. H. El-Sayed, A. Samarkina, J. Xu, J. D. Aquino, M. Xiao, M. Herlyn, I. Yeh, A Technique for the Generation of Mucosal and Acral Melanoma Cell Lines. Pigment Cell & Melanoma Research 39, e70117 (2026).

37. R. Poplin, P.-C. Chang, D. Alexander, S. Schwartz, T. Colthurst, A. Ku, D. Newburger, J. Dijamco, N. Nguyen, P. T. Afshar, S. S. Gross, L. Dorfman, C. Y. McLean, M. A. DePristo, A universal SNP and small-indel variant caller using deep neural networks. Nat Biotechnol 36, 983–987 (2018).

38. M. Martin, M. Patterson, S. Garg, S. O. Fischer, N. Pisanti, G. W. Klau, A. Schöenhuth, T. Marschall, WhatsHap: fast and accurate read-based phasing. bioRxiv [Preprint] (2016). 10.1101/085050.

39. T. Rausch, T. Zichner, A. Schlattl, A. M. Stütz, V. Benes, J. O. Korbel, DELLY: structural variant discovery by integrated paired-end and split-read analysis. Bioinformatics 28, i333–i339 (2012).

40. F. Ramírez, D. P. Ryan, B. Grüning, V. Bhardwaj, F. Kilpert, A. S. Richter, S. Heyne, F. Dündar, T. Manke, deepTools2: a next generation web server for deep-sequencing data analysis. Nucleic Acids Res 44, W160–W165 (2016).

41. Y. Zhang, T. Liu, C. A. Meyer, J. Eeckhoute, D. S. Johnson, B. E. Bernstein, C. Nusbaum, R. M. Myers, M. Brown, W. Li, X. S. Liu, Model-based Analysis of ChIP-Seq (MACS). Genome Biology 9, R137 (2008).

42. M. P. Meers, D. Tenenbaum, S. Henikoff, Peak calling by Sparse Enrichment Analysis for CUT&RUN chromatin profiling. Epigenetics & Chromatin 12, 42 (2019).

43. K. Mendelson, T. C. Martin, C. B. Nguyen, M. Hsu, J. Xu, C. Lang, R. Dummer, Y. Saenger, J. L. Messina, V. K. Sondak, G. Desman, D. Hasson, E. Bernstein, R. E. Parsons, J. T. Celebi, Differential histone acetylation and super-enhancer regulation underlie melanoma cell dedifferentiation. JCI Insight 9 (2024).

44. M. Takahashi, H. B. Chong, S. Zhang, T.-Y. Yang, M. J. Lazarov, S. Harry, M. Maynard, B. Hilbert, R. D. White, H. E. Murrey, C.-C. Tsou, K. Vordermark, J. Assaad, M. Gohar, B. R. Dürr, M. Richter, H. Patel, G. Kryukov, N. Brooijmans, A. S. O. Alghali, K. Rubio, A. Villanueva, J. Zhang, M. Ge, F. Makram, H. Griesshaber, D. Harrison, A.-S. Koglin, S. Ojeda, B. Karakyriakou, A. Healy, G. Popoola, I. Rachmin, N. Khandelwal, J. R. Neil, P.-C. Tien, N. Chen, T. Hosp, S. van den Ouweland, T. Hara, L. Bussema, R. Dong, L. Shi, M. Q. Rasmussen, A. C. Domingues, A. Lawless, J. Fang, S. Yoda, L. P. Nguyen, S. M. Reeves, F. N. Wakefield, A. Acker, S. E. Clark, T. Dubash, J. Kastanos, E. Oh, D. E. Fisher, S. Maheswaran, D. A. Haber, G. M. Boland, M. Sade-Feldman, R. W. Jenkins, A. N. Hata, N. M. Bardeesy, M. L. Suvà, B. R. Martin, B. B. Liau, C. J. Ott, M. N. Rivera, M. S. Lawrence, L. Bar-Peled, DrugMap: A quantitative pan-cancer analysis of cysteine ligandability. Cell 187, 2536–2556.e30 (2024).

45. Open2C, N. Abdennur, G. Fudenberg, I. M. Flyamer, A. A. Galitsyna, A. Goloborodko, M. Imakaev, S. V. Venev, Pairtools: From sequencing data to chromosome contacts. PLOS Computational Biology 20, e1012164 (2024).

46. X. Wang, J. Xu, B. Zhang, Y. Hou, F. Song, H. Lyu, F. Yue, Genome-wide detection of enhancer-hijacking events from chromatin interaction data in rearranged genomes. Nat Methods 18, 661–668 (2021).

47. X. Wang, Y. Luan, F. Yue, EagleC: A deep-learning framework for detecting a full range of structural variations from bulk and single-cell contact maps. Sci Adv 8, eabn9215.

48. M. E. Fornace, J. Huang, C. T. Newman, A. Nanjundiah, N. J. Porubsky, M. B. Pierce, N. A. Pierce, NUPACK: Computational Nucleic Acid Analysis and Design. ACS Synth. Biol. 15, 1426–1441 (2026).

49. A. P. Heath, V. Ferretti, S. Agrawal, M. An, J. C. Angelakos, R. Arya, R. Bajari, B. Baqar, J. H. B. Barnowski, J. Burt, A. Catton, B. F. Chan, F. Chu, K. Cullion, T. Davidsen, P.-M. Do, C. Dompierre, M. L. Ferguson, M. S. Fitzsimons, M. Ford, M. Fukuma, S. Gaheen, G. L. Ganji, T. I. Garcia, S. S. George, D. S. Gerhard, F. Gerthoffert, F. Gomez, K. Han, K. M. Hernandez, B. Issac, R. Jackson, M. A. Jensen, S. Joshi, A. Kadam, A. Khurana, K. M. J. Kim, V. E. Kraft, S. Li, T. M. Lichtenberg, J. Lodato, L. Lolla, P. Martinov, J. A. Mazzone, D. P. Miller, I. Miller, J. S. Miller, K. Miyauchi, M. W. Murphy, T. Nullet, R. O. Ogwara, F. M. Ortuño, J. Pedrosa, P. L. Pham, M. Y. Popov, J. J. Porter, R. Powell, K. Rademacher, C. P. Reid, S. Rich, B. Rogel, H. Sahni, J. H. Savage, K. A. Schmitt, T. J. Simmons, J. Sislow, J. Spring, L. Stein, S. Sullivan, Y. Tang, M. Thiagarajan, H. D. Troyer, C. Wang, Z. Wang, B. L. West, A. Wilmer, S. Wilson, K. Wu, W. P. Wysocki, L. Xiang, J. T. Yamada, L. Yang, C. Yu, C. K. Yung, J. C. Zenklusen, J. Zhang, Z. Zhang, Y. Zhao, A. Zubair, L. M. Staudt, R. L. Grossman, The NCI Genomic Data Commons. Nat Genet 53, 257–262 (2021).

