## Supplementary_Materials for "WRN deficiency generates oncogenic rearrangements at TA-rich tandem repeats"

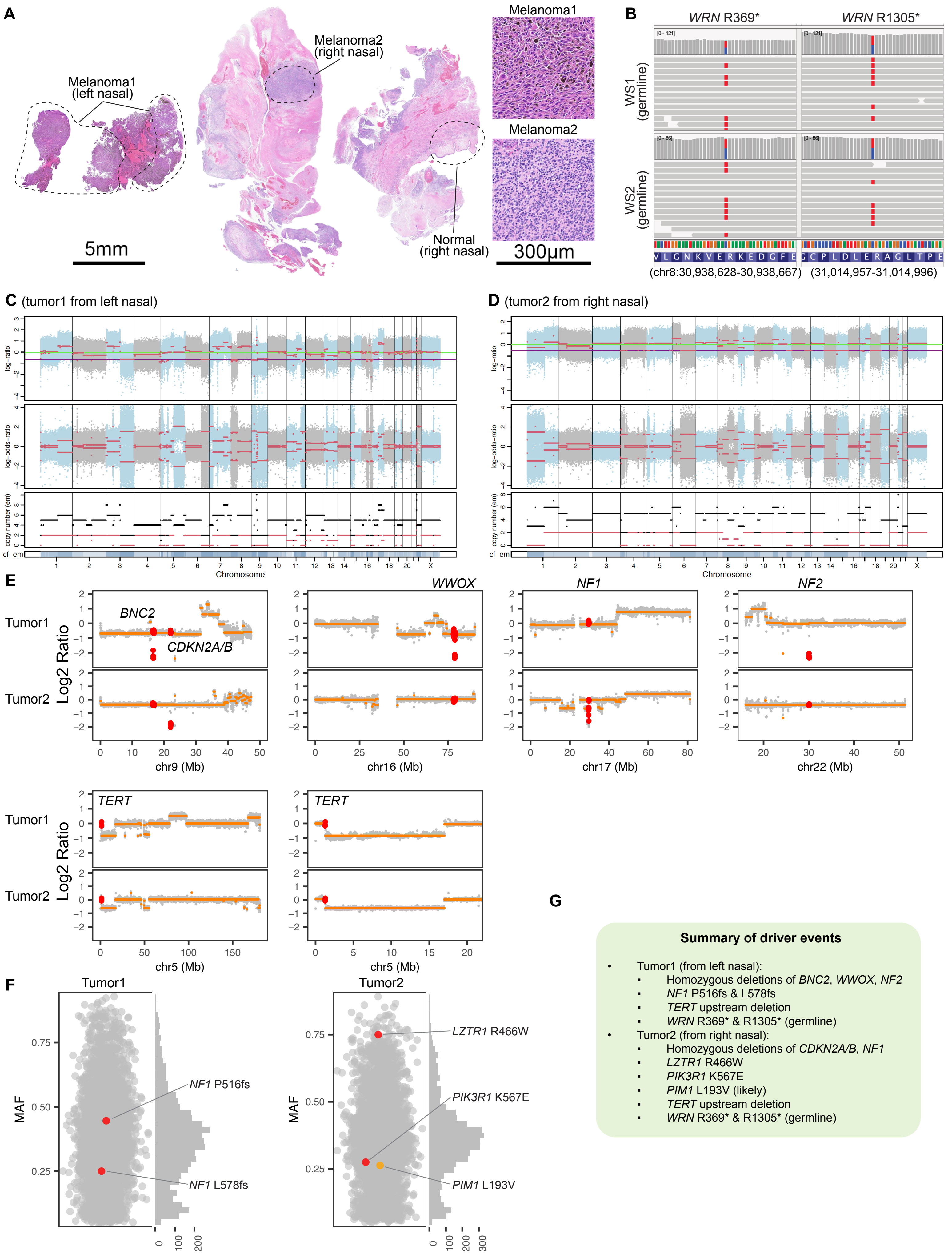
A
Melanoma1 (left nasal)
Melanoma2 (right nasal)
Normal (right nasal)
5mm
Melanoma1
Melanoma2
300µm
B
WRN R369*
WRN R1305*
WS1 (germline)
WS2 (germline)
(chr8:30,938,628-30,938,667)
(31,014,957-31,014,996)
C (tumor1 from left nasal)
D (tumor2 from right nasal)
log−ratio
log−odds−ratio
copy number (em)
cf−em
Chromosome
E
Tumor1
Tumor2
Log2 Ratio
BNC2
CDKN2A/B
WWOX
NF1
NF2
chr9 (Mb)
chr16 (Mb)
chr17 (Mb)
chr22 (Mb)
TERT
chr5 (Mb)
F
Tumor1
Tumor2
MAF
NF1 P516fs
NF1 L578fs
LZTR1 R466W
PIK3R1 K567E
PIM1 L193V
G
Summary of driver events
• Tumor1 (from left nasal):
▪ Homozygous deletions of BNC2, WWOX, NF2
▪ NF1 P516fs & L578fs
▪ TERT upstream deletion
▪ WRN R369* & R1305* (germline)
• Tumor2 (from right nasal):
▪ Homozygous deletions of CDKN2A/B, NF1
▪ LZTR1 R466W
▪ PIK3R1 K567E
▪ PIM1 L193V (likely)
▪ TERT upstream deletion
▪ WRN R369* & R1305* (germline)

**Figure S1. Genetic summary of two primary melanomas from patient WS1. A.** Hematoxylin and eosin-stained sections of tumors from the left and right nasal cavity, respectively. The normal and tumor areas were microdissected as indicated. **B.** IGV screenshots showing two heterozygous *WRN* nonsense mutations, R369* and R1305*, in the germline. **C** and **D.** Copy number profile of the two melanomas. The upper panels show $\log_2$ ratio values; the middle panel shows log-odds ratios reflecting allelic imbalance of germline heterozygous variants in tumor versus germline; and the lower panel shows estimated absolute copy numbers across the genome. **E.** Magnified copy-number plots highlighting alterations in selected genes. Specifically, *BNC2*, *WWOX* and *NF2* were homozygously deleted in melanoma 1, whereas *CDKN2A/B*, and *NF1* were homozygously deleted in melanoma 2. Both tumors share the same 15.7Mb deletion on chromosome 5p near *TERT*. **F.** Scatterplots showing the mutant allele frequency (MAF) of somatic mutations. **G.** Summary of driver events in the two melanomas.

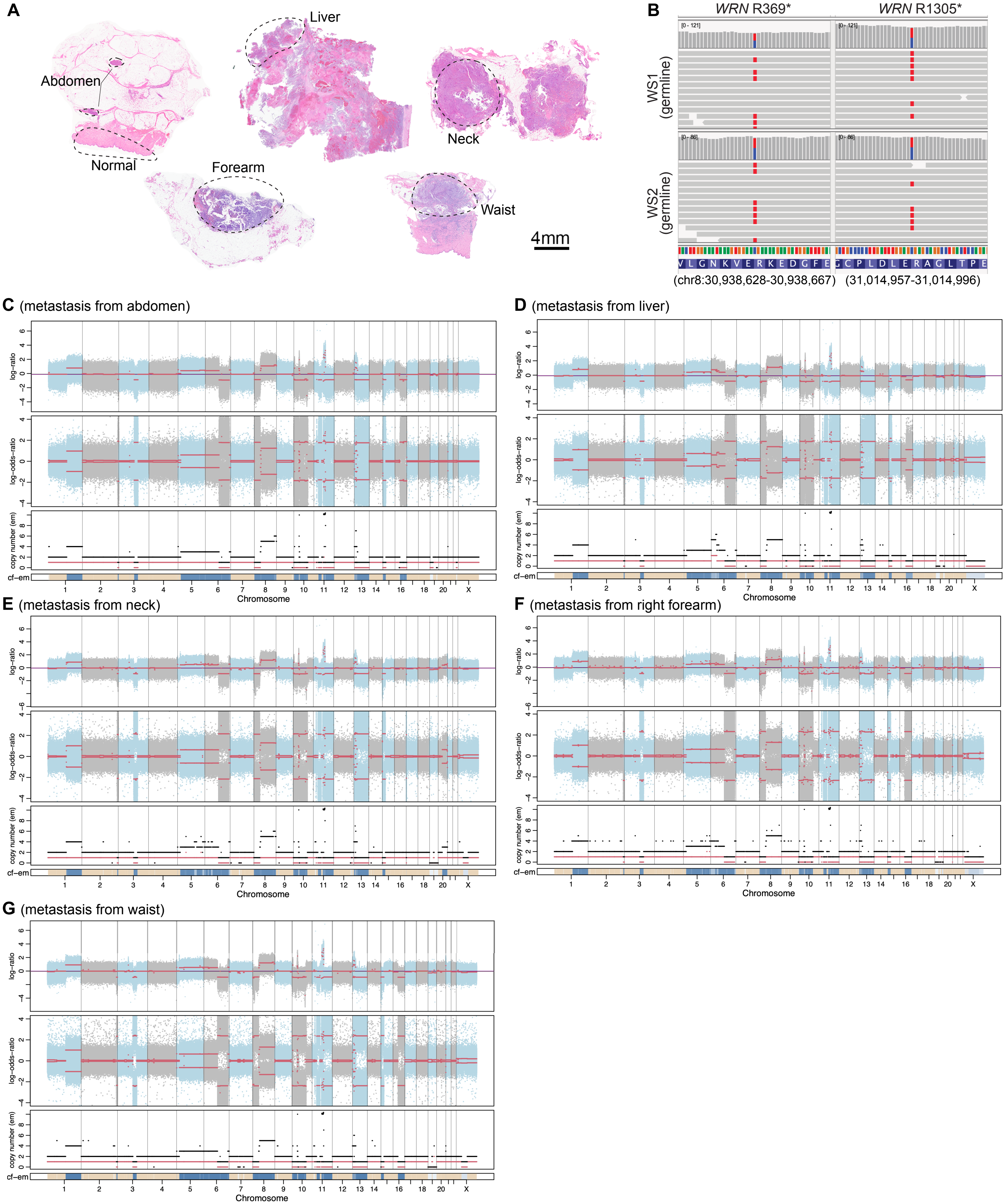
A
Liver
Abdomen
Neck
Normal
Forearm
Waist
4mm
B
WRN R369*
WRN R1305*
WS1 (germline)
WS2 (germline)
[0 - 121]
[0 - 86]
V L G N K V E R K E D G F E
G C P L D L E R A G L T P E
(chr8:30,938,628-30,938,667)
(31,014,957-31,014,996)
C (metastasis from abdomen)
D (metastasis from liver)
E (metastasis from neck)
F (metastasis from right forearm)
G (metastasis from waist)
log−ratio
log−odds−ratio
copy number (em)
cf−em
Chromosome
1
2
3
4
5
6
7
8
9
10
11
12
13
14
16
18
20
X

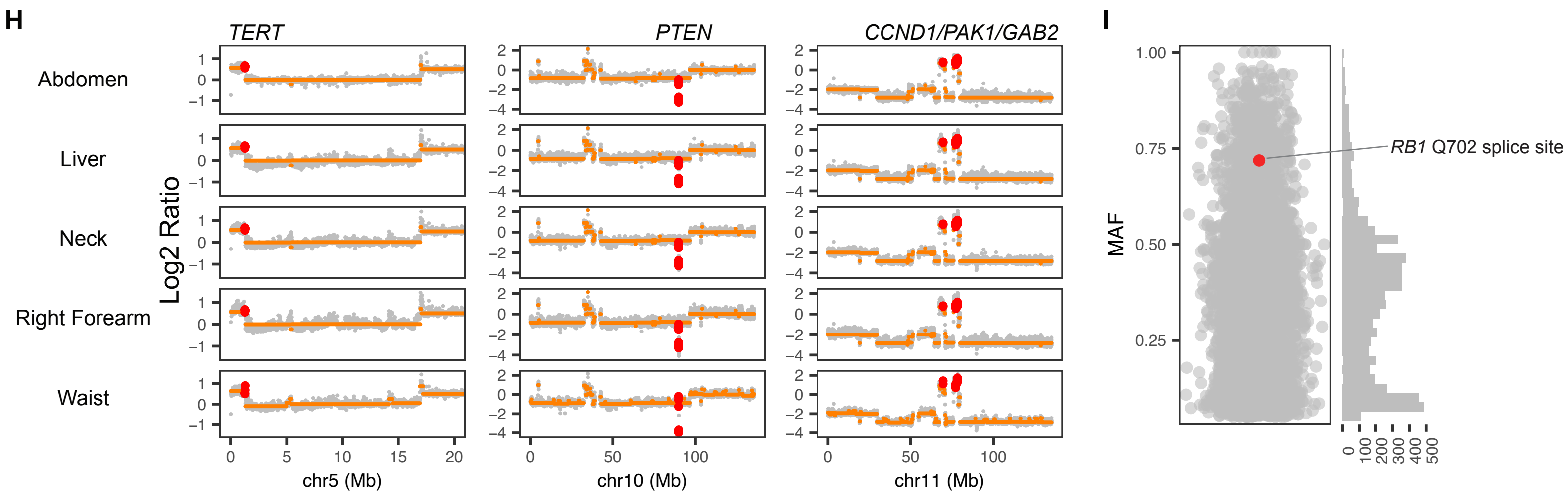


**Figure S2. Genetic summary of five metastatic lesions from patient WS2. A.** Hematoxylin and eosin-stained sections of metastases from abdomen, liver, neck, forearm and waist, respectively. The normal and tumor areas were microdissected as indicated. **B.** IGV screenshots showing two heterozygous WRN nonsense mutations, R369* and R1305*, in the germline. **C, D, E, F** and **G.** Copy number profile of the two melanomas. The upper panels show log2 ratio values; the middle panel shows log-odds ratios reflecting allelic imbalance of germline heterozygous variants in tumor versus germline; and the lower panel shows estimated absolute copy numbers across the genome. **H.** Magnified copy-number plots highlighting alterations in selected genes. Specifically, all metastases share the same 15.7Mb deletion on chromosome 5p near *TERT*, *PTEN* homozygous deletion, and amplifications of *CCND1*, *PAK1* and *GAB2*. **I.** Scatterplots showing the mutant allele frequency (MAF) of somatic mutations, with the abdomen metastasis as an example. *RB1* Q702 splice site mutation is shared in all metastases.

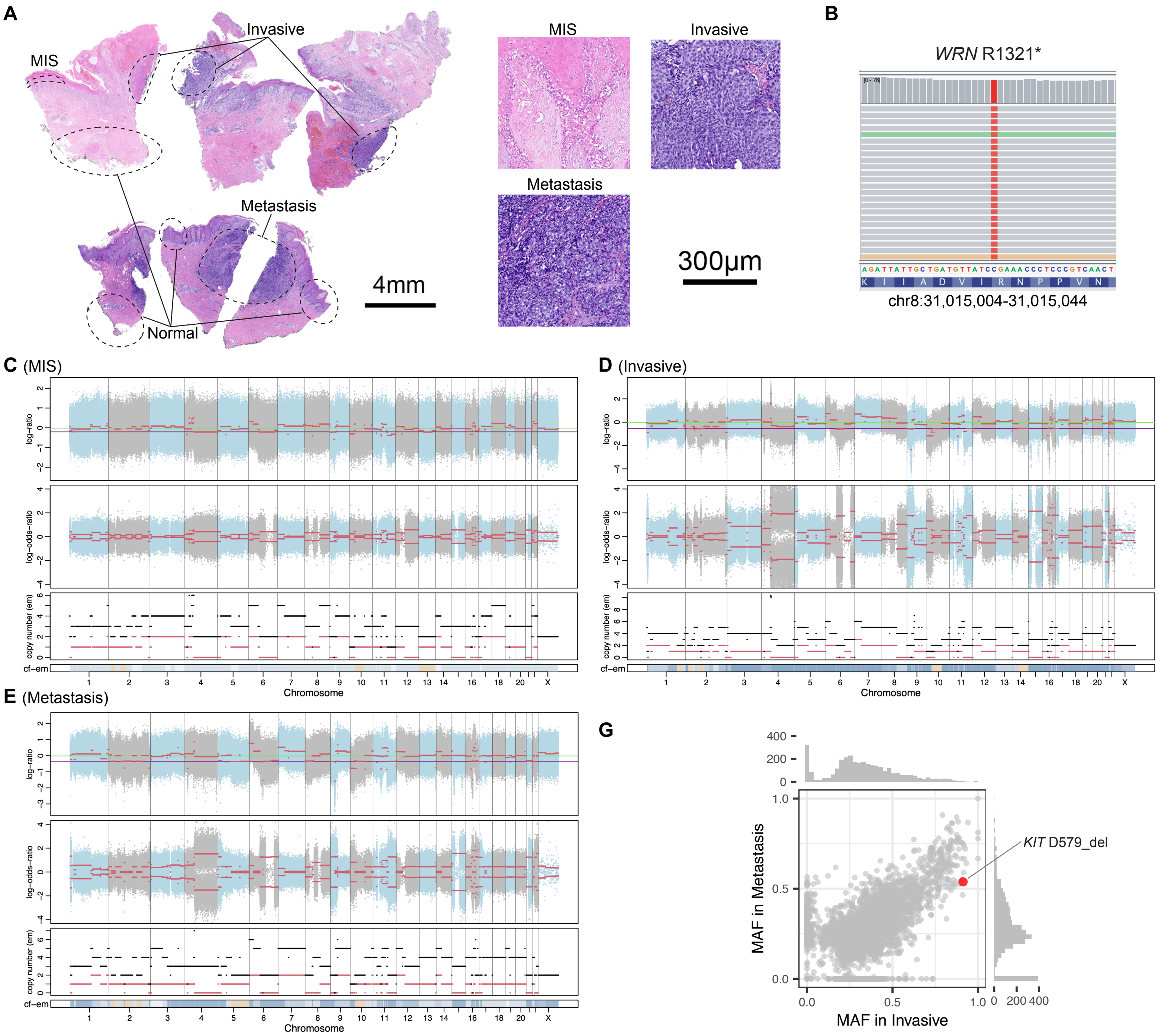


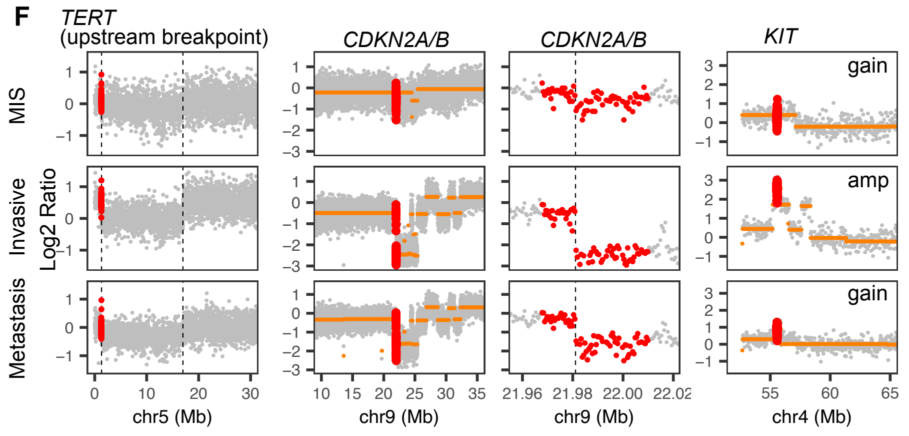


**Figure S3. Genetic summary of the acral melanoma from patient WS3. A.** Hematoxylin and eosin-stained sections of melanoma in situ (MIS) and invasive from right heel, and local metastasis from right medial ankle. The normal and tumor areas were microdissected as indicated. **B.** IGV screenshots showing one homozygous *WRN* nonsense mutations, R1321*, in the germline. **C, D,** and **E.** Copy number profile of the three tumor areas. The upper panels show log2 ratio values; the middle panel shows log-odds ratios reflecting allelic imbalance of germline heterozygous variants in tumor versus germline; and the lower panel shows estimated absolute copy numbers across the genome. **F.** Magnified copy-number plots highlighting alterations in selected genes. Specifically, all metastases share the same 15.7Mb deletion on chromosome 5p near *TERT* and *CDKN2A* homozygous deletion. *KIT* was gained in MIS and metastasis but highly amplified in invasive. **G.** Scatterplots showing the mutant allele frequency (MAF) of somatic mutations in invasive and metastasis. a putative driver mutation, *KIT* D579del, was observed in both areas. Its increased MAF in the invasive lesion corresponds to *KIT* amplification in this area.

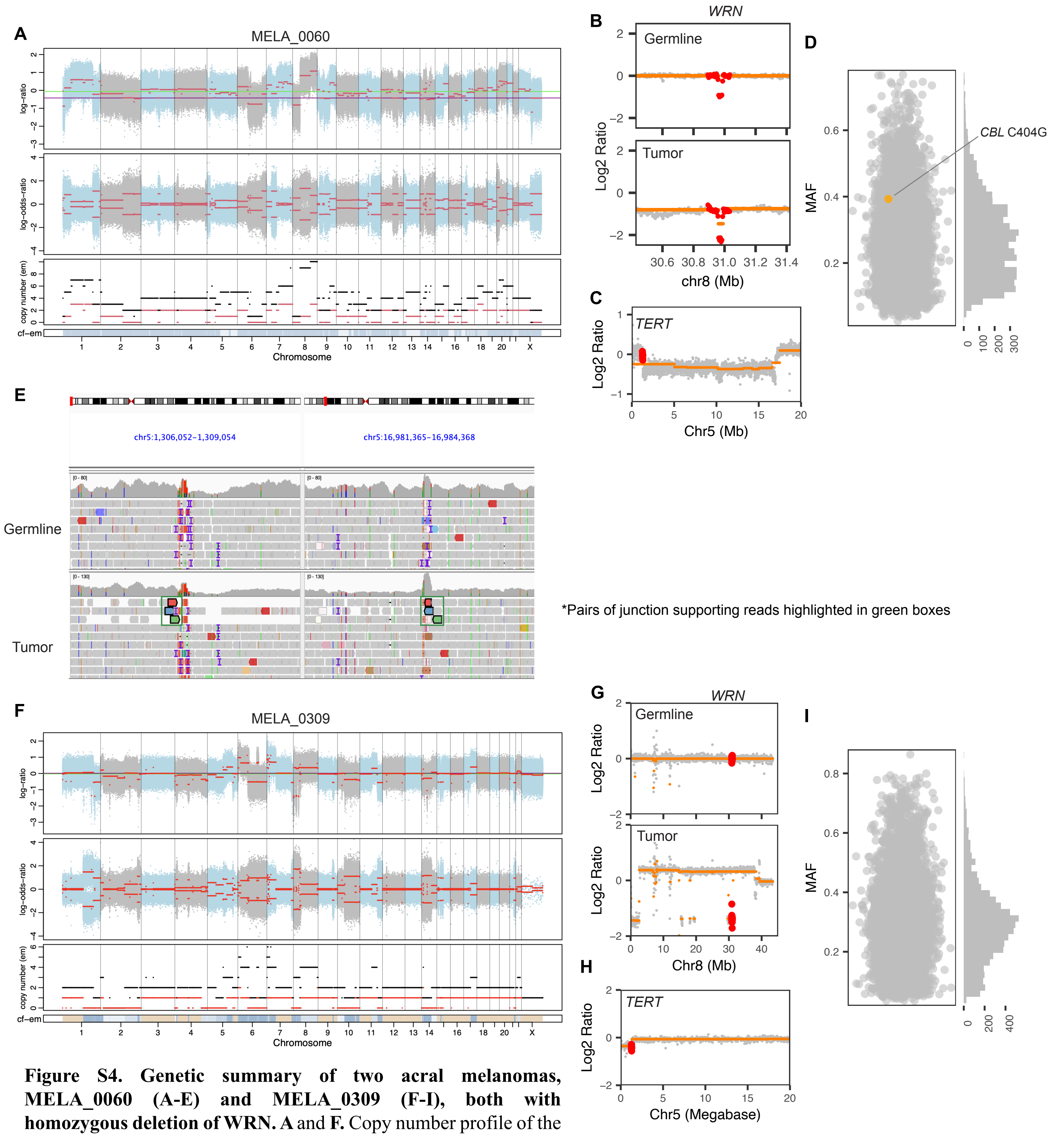


**Figure S4. Genetic summary of two acral melanomas, MELA_0060 (A-E) and MELA_0309 (F-I), both with homozygous deletion of WRN. A** and **F.** Copy number profile of the two tumors. The upper panels show log2 ratio values; the middle panel shows log-odds ratios reflecting allelic imbalance of germline heterozygous variants in tumor versus germline; and the lower panel shows estimated absolute copy numbers across the genome. **B** and **G.** Magnified copy-number plots of WRN in both germline and tumor for the two patients. Both tumors show homozygous deletion of WRN. For MELA_0060, a heterozygous deletion spanning exons 19-23 is observed in the germline. **C** and **H.** Magnified copy-number plots of chromosome 5p. The 15.7Mb deletion on chromosome 5p was observed only in MELA_0060. **D** and **I.** Scatterplots showing the mutant allele frequency (MAF) of somatic mutations. A likely driver mutation CBL C404G was noted for MELA_0060. **E.** IGV screenshot showing pairs of reads that support the direct connection of the two TA-repeat breakpoints ($BP_{tel}$ and $BP_{cen}$) in MELA_0060.

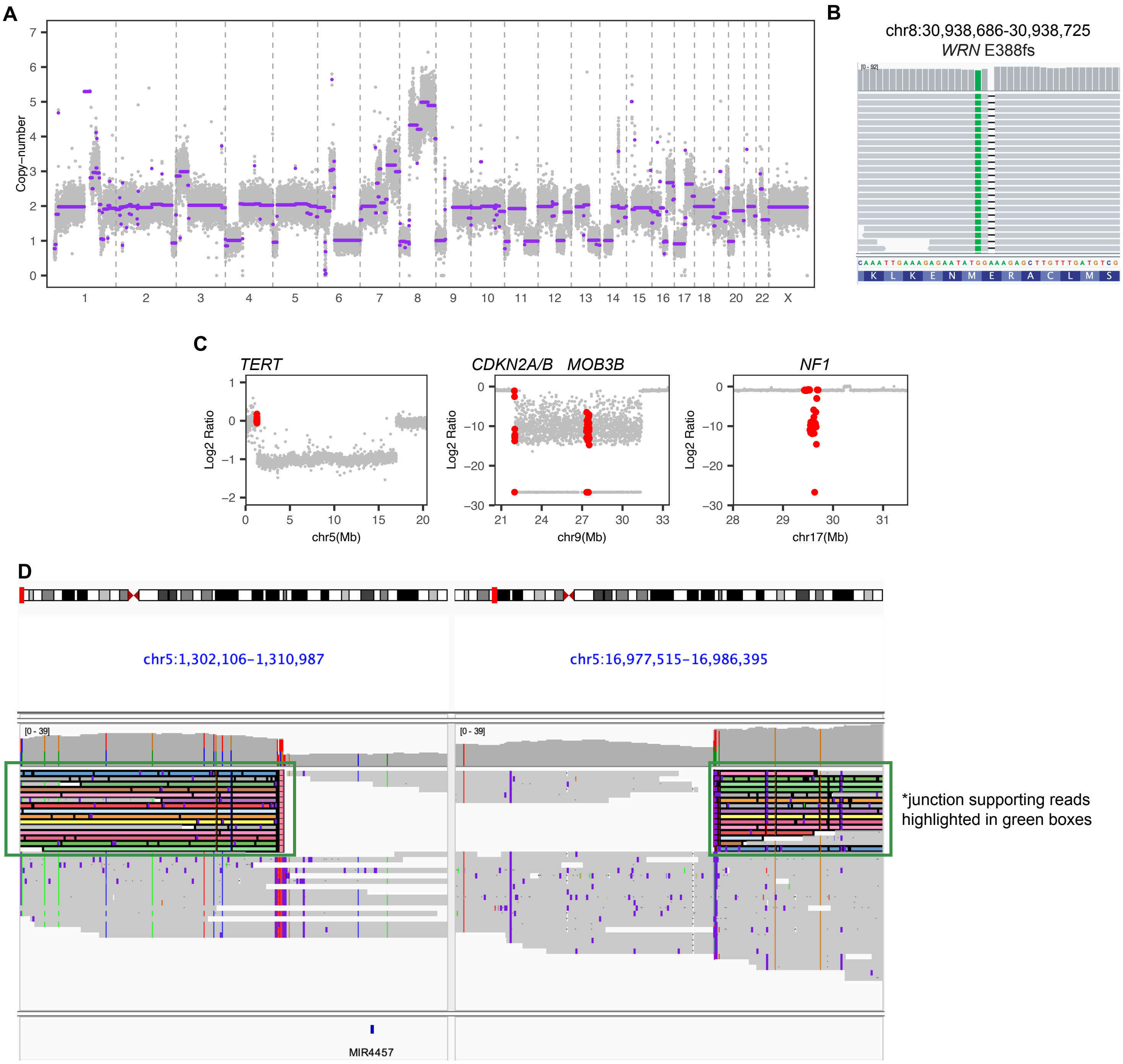


**Figure S5. Genetic summary of the mucosal melanoma cell line MTB3004A1S. A.** The copy number profile across the genome. **B.** IGV screenshots showing hemizygous *WRN* mutations M387I and E388fs. **C.** Magnified copy-number plots highlighting alterations of selected genes, including the 15.7Mb deletion on chromosome 5p near *TERT*, and the homozygous deletions of *CDKN2A*, *MOB3B* and *NF1*. **D.** IGV screenshot showing long reads that support the direct connection of the two TA-repeat breakpoints ($BP_{tel}$ and $BP_{cen}$) in MELA_0060.

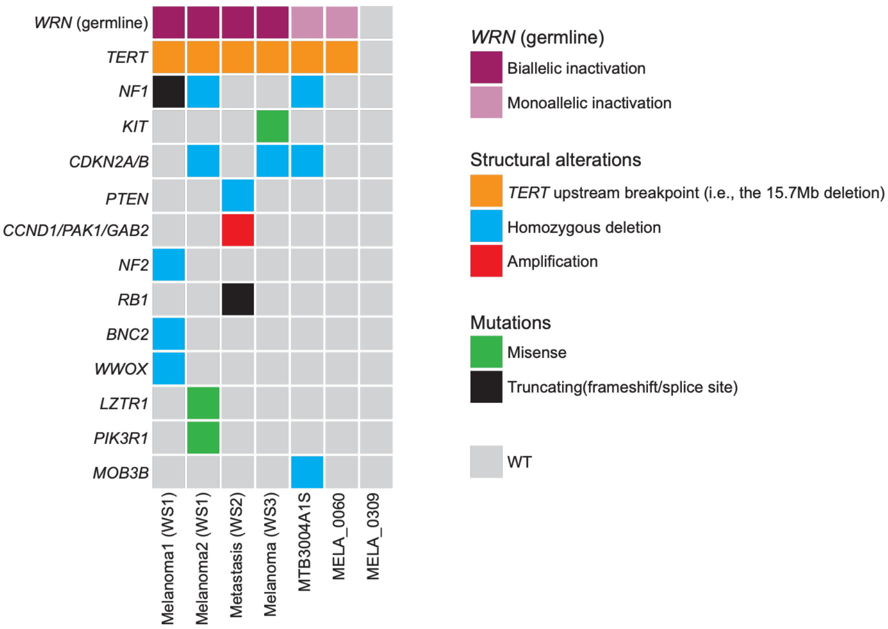


**Figure S6. Oncoplot showing the driver alterations of the 7 WRN-null melanomas.** The germline status of WRN is also summarized.

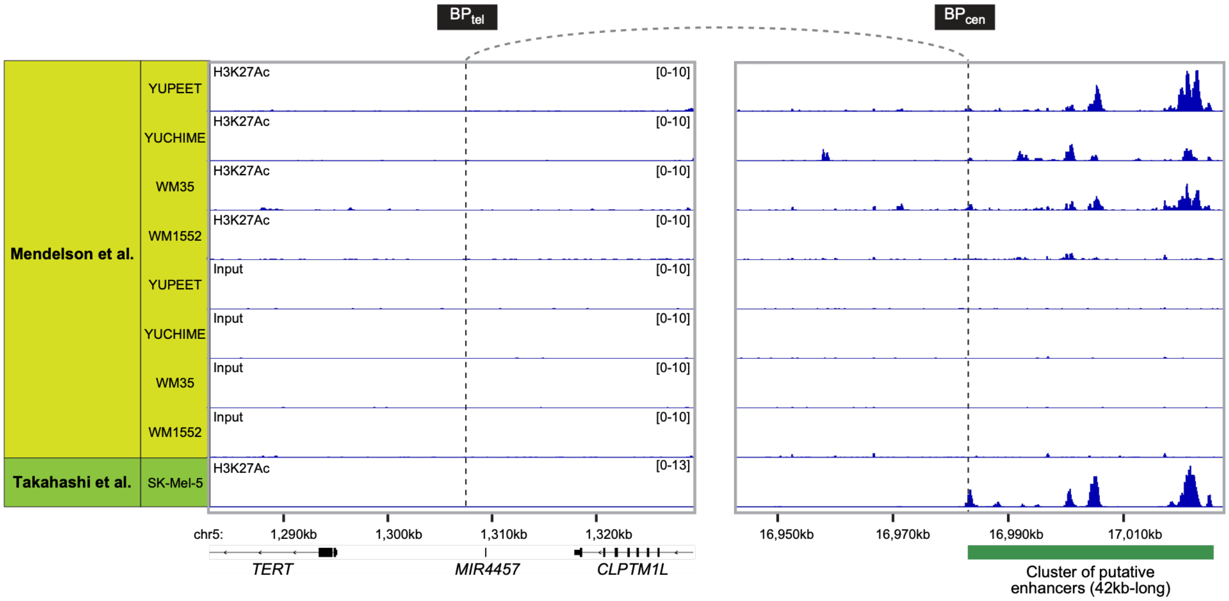


**Figure S7. H3K27ac ChIP-seq signals of regions surrounding $BP_{tel}$ and $BP_{cen}$ in melanoma cell lines from two previous studies.** Note that all the cell lines were derived from cutaneous melanomas**.** The green box indicates the cluster of enhancer regions identified in MTB3004A1S (**Fig. 2**).

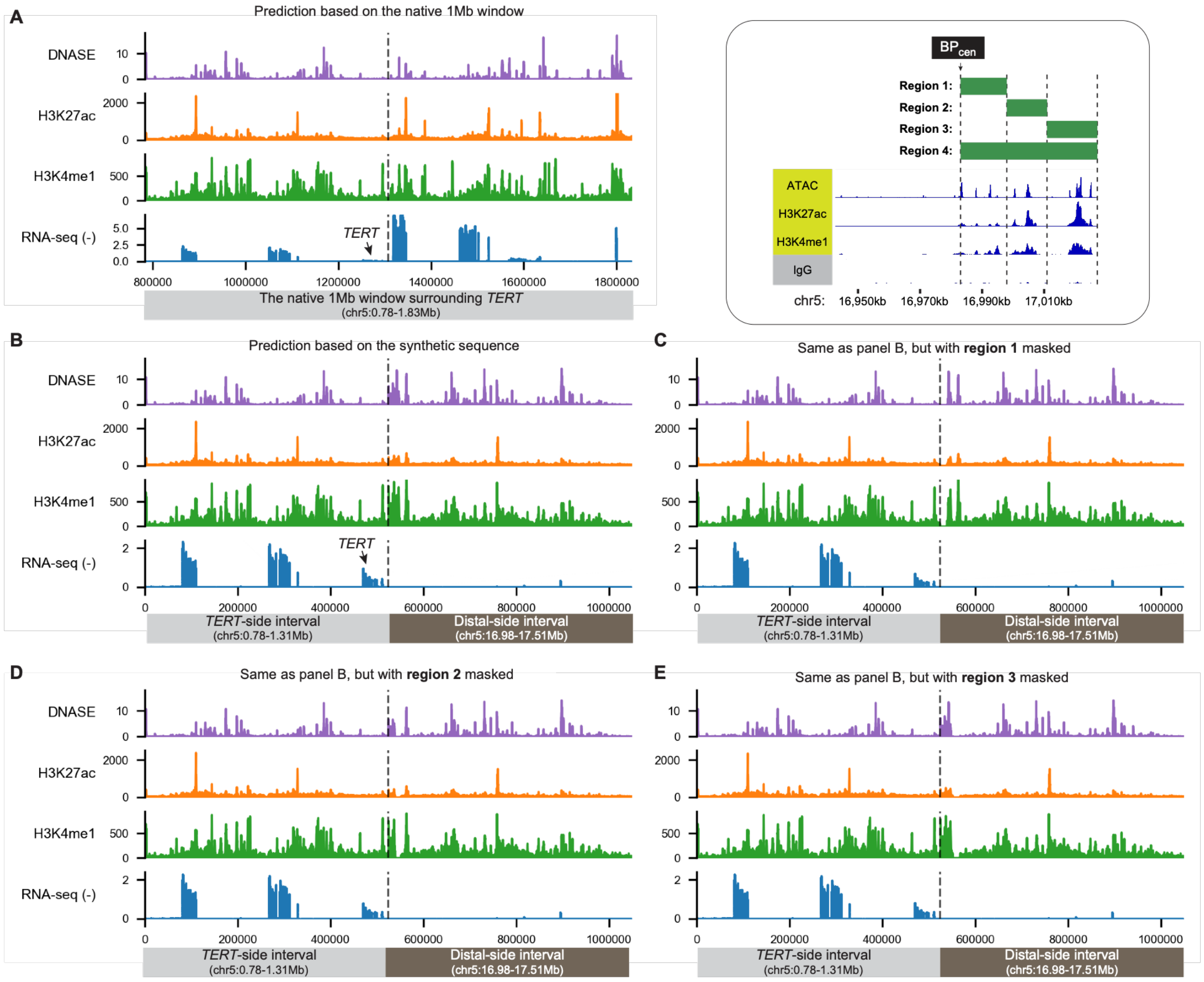


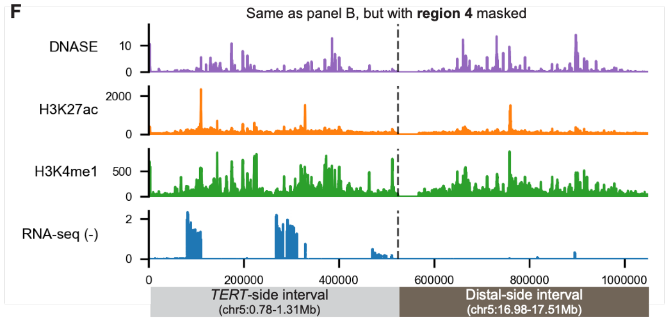


G

| | TERT expression (intensity per bps) | Fold change (relative to panel B) |
|---|---|---|
| Native sequence (panel A) | 0.058 | 0.153 |
| Synthetic sequence (panel B) | 0.380 | 1 |
| Synthetic sequence, with region 1 removed (panel C) | 0.249 | 0.655 |
| Synthetic sequence, with region 2 removed (panel D) | 0.311 | 0.817 |
| Synthetic sequence, with region 3 removed (panel E) | 0.313 | 0.823 |
| Synthetic sequence, with region 4 removed (panel F) | 0.167 | 0.439 |

**Figure S8. AlphaGenome prediction of *TERT* expression, DNase I hypersensitivity, and histone mark profiles in the melanocytic lineage. (A)** Epigenomic and transcriptional predictions for the native 1Mb reference genomic window surrounding the *TERT* locus. **(B)** Computational prediction for the 1Mb synthetic rearranged sequence generated by a fusion of equal halves flanking $BP_{tel}$ and $BP_{cen}$, modeling the genomic topology following the somatic deletion event. **(C–F)** Predictive profiles for the synthetic sequence variants following localized in silico sequence masking, where specific targeted intervals are replaced with non-informative "N" sequences. **(G)** Summary of quantitative *TERT* expression levels across all simulated configurations and their respective fold-changes relative to the baseline synthetic rearranged model depicted in **(B)**. The vertical dashed lines denote the native position of $BP_{tel}$ in **(A)**, and the novel structural junction point merging $BP_{tel}$ and $BP_{cen}$ in **(B–F)**. All predictions were performed based on the foreskin melanocyte primary cells (Cell Ontology: CL2000045). The genomic coordinates of both panels are based on hg38.

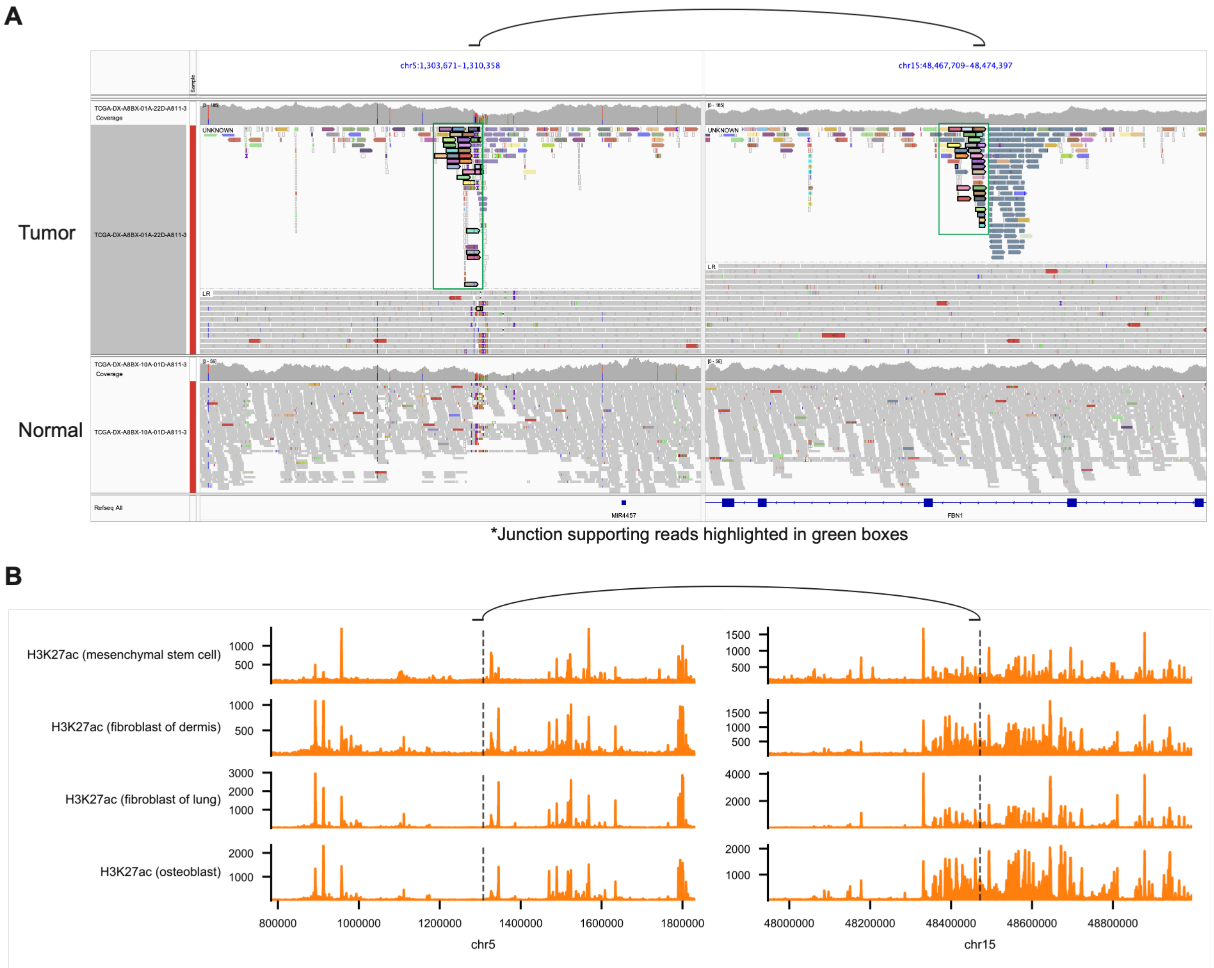


**Figure S9. The *TERT* locus is rearranged to chromosome 15 in sample DX-A8BX. (A)** IGV screenshots showing detailed read mapping at $BP_{tel}$ and its partner breakpoint on chromosome 15q. In the tumor sample, reads were grouped by pair orientation into unknown orientation reads (upper track, consisting predominantly of junction-supporting reads) and conventional left-right (LR) orientation reads. The read mapping indicates that the chromosome 15q locus is involved in multiple rearrangements. The arch at the top indicates the rearrangement involving *TERT*. The reads in normal are squished. **(B)** H3K27ac profiles surrounding the two breakpoints in mesenchymal stem cells, fibroblasts, and osteoblast cell lines based on AlphaGenome predictions. The genomic coordinates of both panels are based on hg38.
